# Precision-Weighted Updating Explains Serial Dependence Across Sensory and Contextual Transitions

**DOI:** 10.64898/2026.06.06.730048

**Authors:** Chunyu Qu, Zhuanghua Shi

## Abstract

Serial dependence is influenced by sensory uncertainty and by the continuity of context, but it remains controversial whether these influences reflect separate mechanisms or different expressions of a shared updating process. Across two time reproduction experiments (N = 44), we examined how motion coherence and transitions between coherence levels modulated the attraction toward the preceding duration, while controlling for central tendency. In Experiment 1, low current coherence led to stronger serial dependence than high coherence, whereas central tendency did not change. In Experiment 2, in which abrupt coherence onsets and a color change marked each encoding interval, serial dependence was elevated when low coherence repeated across trials. A three-state Kalman filter, comprising a fast duration estimate, a slowly learned mean, and a slow response bias, captured these patterns. In the fitted models, coherence did not change how strongly the current duration was weighted. It changed how much of the trace left by the preceding trial was retained in the prior for the current estimate, and in Experiment 2 a change of coherence interrupted this retention. With all parameters fixed, the same model predicted how both biases change when the volatility of the stimulus sequence is manipulated, in the data of Glasauer and Shi (2022). These findings support a precision-weighted updating account in which current evidence quality and perceptual continuity regulate the use of recent history.

## 1 INTRODUCTION

When an observer reproduces a brief interval, the response depends on more than the interval just presented. The interval of the preceding trial leaves a trace, and the current reproduction is drawn toward it. Such traces are useful when successive events resemble each other, because the recent past then carries information about the present. They become a source of error when the present event is unrelated to the one before. This is a general stability–flexibility problem for cognition: the mind must keep recent information active long enough to support perception and action, yet limit its influence when that information no longer applies.

This tension is central to history-dependent cognition: the pervasive influence of recent experience on current perception, attention, memory, and action (Kiyonaga et al., 2017; Wang et al., 2025; Wu et al., 2026). Such dependence is often adaptive. In a world where many properties persist over short intervals, relying on the recent past can reduce noise and support continuity. Yet persistence is not guaranteed. When the environment changes, the same history dependence can bias judgment away from the current state of the world; a large-scale mega-analysis found that serial dependence increased perceptual decision error across its compiled data sets (Ozkirli et al., 2026). A cognitive system must therefore regulate how strongly prior information influences present estimates, increasing stability when continuity is likely and reducing carryover when change is likely.

Two well-established biases provide measurable signatures of this regulation. Serial dependence refers to the attraction of current judgments toward recently encountered stimuli, as when the perceived orientation, or motion direction is pulled toward the preceding trial (J. Fischer & Whitney, 2014). Central tendency refers to regression toward the long-run mean of the stimulus distribution, a classic phenomenon in judgment and time perception (Hollingworth, 1910; Vierordt, 1868). Although these effects are often discussed separately, both can be viewed as forms of temporal integration over different timescales: observers combine current evidence with information accumulated from recent and longer-term experience. Recent computational accounts have emphasized this common structure, showing how sequential biases can arise from iterative estimation processes that balance incoming sensory evidence against prior expectations (Cicchini et al., 2018; Glasauer & Shi, 2022; Shi et al., 2013).

One central determinant of this balance is precision. When current sensory evidence is reliable, it should dominate the estimate; when it is noisy or uncertain, prior information should receive more weight. This principle follows from optimal cue integration, Bayesian perception, and predictive-processing accounts, in which sensory evidence and prior expectations are weighted by their relative reliability (Ernst & Banks, 2002; Feldman & Friston, 2010; van Bergen & Jehee, 2019). Consistent with this view, several studies have found stronger serial dependence when the current stimulus is uncertain, and duration judgments under memory pressure show stronger regression toward stable prior information (Ceylan et al., 2021; Markov et al., 2024; Zang et al., 2026). Together, these findings suggest that the pull of the past is not fixed, but varies with the reliability of current evidence.

Yet how much a preceding event contributes may depend not only on stimulus reliability, but also on whether the present event continues it. Serial dependence is tuned to the similarity of successive stimuli in space, time, and features, which has been taken to indicate that it supports the continuity of perceived objects (Manassi & Whitney, 2024; Pascucci et al., 2023), and the proposal of spatiotemporal routines likewise emphasizes how spatial and temporal context organize perception and behavior (Pascucci & Kristjánsson, 2026). Similar boundaries structure perception and memory more generally. Event segmentation theories propose that observers parse continuous experience into meaningful units, with boundaries marking moments when predictions become less reliable (Zacks & Swallow, 2007). Such boundaries alter temporal binding and memory organization, separating information within an event from information across events (Baldassano et al., 2017; DuBrow & Davachi, 2016). In serial dependence, task relevance and contextual continuity likewise influence whether information from the previous trial carries forward. Merely encoding a previous target may be insufficient if that feature was not task relevant, and switches in task demands or feature relevance can weaken attractive carryover or reveal opposing biases (Bae & Luck, 2020; Bliss et al., 2017; Ceylan & Pascucci, 2023; Cheng, Chen, Glasauer, et al., 2024; Cheng, Chen, & Shi, 2024; Cheng, Chen, Yang, et al., 2024). These observations raise the possibility that a perceptual boundary between successive events limits how much of the preceding trace enters the current estimate.

The relation between sensory uncertainty and contextual continuity on serial dependence remains unresolved. These factors are often treated as distinct modulators: stimulus quality affects how strongly the past is weighted, whereas task context affects whether past information is selected, gated, or discounted. However, both may reflect a common inferential problem: estimating whether previous information remains predictive of the present. A further challenge comes from an asymmetry in reliability-related effects. Simple direct integration accounts predict that uncertainty on both the current and previous trials should influence the strength of serial dependence, because the final estimate combines information from both. In several recent paradigms, however, serial dependence has appeared to depend more strongly on current trial uncertainty than on previous trial uncertainty (Ceylan et al., 2021; Gallagher & Benton, 2022), including across features (You et al., 2023). Gallagher and Benton (2022) described this pattern as “seemingly anti-Bayesian,” because it is difficult to reconcile with a static weighted average of current and previous observations. A dynamic state-space account (e.g., Glasauer & Shi, 2022) offers one way to reconcile this pattern: what the previous trial contributes is already contained in the observer’s state when the current stimulus arrives, whereas the current trial determines how that state is combined with the new evidence.

The present study examines these issues in two time reproduction experiments. The intervals were presented as random-dot motion whose coherence, either low or high, varied randomly from trial to trial as a manipulation of stimulus quality. A trial therefore repeated or switched the coherence of the preceding trial with equal probability, so that successive trials differed in perceptual continuity whereas the statistics of the sequence were constant. The onset and offset of the interval were gradual in Experiment 1 and sharply marked in Experiment 2, which set each interval apart as a separate perceptual event. Three accounts make different predictions for this design. Static reliability weighting predicts that serial dependence varies with the coherence of both the previous and the current trial, whereas an account in which only current uncertainty matters predicts that it varies with current coherence alone. An account in which the trace is carried forward only while the perceptual context continues predicts stronger serial dependence when coherence repeats than when it switches. Based on the literature, we expected that current coherence would influence serial dependence, and that the repetition of coherence would matter more when the intervals were sharply marked (Experiment 2). To determine whether stimulus quality and continuity act through distinct mechanisms or through one updating process, we used a three-state Kalman filter that tracks the current duration, the learned mean of the durations, and a slow response bias. We compared model variants that differ in where coherence and a switch of coherence act. Finally, to test the generalizability of the model, we applied the same model to published data in which the volatility of the duration sequence was manipulated (Glasauer & Shi, 2022).

## 2 METHOD

### 2.1 Participants

Forty-four participants completed the two experiments (Experiment 1: *N* = 22, 11 males, 11 females, age 19-24, *M* = 21.23, *SD* = 1.45; Experiment 2: *N* = 22, 12 males, 10 females, age 24-35, *M* = 29.14, *SD* = 3.00). All participants were recruited through Prolific and completed the experiment remotely via Pavlovia. Participants confirmed normal or corrected-to-normal vision and right-handedness. The a priori power analysis targeted the basic within-participant serial dependence effect (two-sided, *d*z = 0.70, alpha = .05, power = .80), based on effect sizes in prior studies (Bae & Luck, 2020; J. Fischer & Whitney, 2014), and indicated a minimum sample size of 19 (Faul et al., 2007). With *N* = 22 per experiment, the design has 80% power for within-participant effects of *d*z ≥ 0.63 (Appendix A). Participants were included in the analyzed sample only if more than 80% of trials remained after preprocessing; retention was defined as the proportion of trials with valid responses (i.e., not missing and not excessively short or long in reproduction duration). All participants gave informed consent and received £9 per hour. The study was approved by the Ethics Committee of the Department of Psychology at LMU Munich (date: 06.04.2022).

### 2.2 Stimuli and apparatus

Both experiments were programmed in PsychoJS (v2023.1.3) and presented via Pavlovia. Stimuli consisted of random-dot kinematograms (RDKs) comprising 360 green dots (approximately 0.03° in diameter) moving within a circular aperture of 3.5° radius at a speed of 1.75°/s, updated at 60 Hz. To account for differences in screen size across participants’ devices, a credit-card calibration procedure was used: participants held a standard credit card against the screen and adjusted an on-screen rectangle to match its dimensions, providing a pixels-per-centimeter conversion factor that scaled all stimuli to physical units. Visual angles were then computed assuming a nominal viewing distance of 57 cm.

### 2.3 Procedure

Both experiments used a time reproduction paradigm (Figure 1A). Each trial began with a 500-ms random motion pre-mask to eliminate residual motion aftereffects. Coherent motion then appeared at one of two coherence levels (30% or 70%) for the target duration. After encoding, a 500-ms random motion post-mask was presented to eliminate residual motion signals, followed by a central letter “T” for 500 ms to signal reproduction onset. Participants pressed and held the ↓ key to reproduce the perceived interval; while the key was held, the RDK reappeared with the same motion direction, stopping when the key was released. The press duration was recorded as the reproduced interval (maximum 6 s). During practice, feedback on reproduction accuracy was given by filling one of five horizontally arranged circles according to the relative error ratio: the leftmost (error ratio < –30%, red), the second from the left (error ratio between –30% and –5%, pink), the middle (error ratio between –5% and +5%, green), the second from the right (error ratio between +5% and +30%, pink), and the rightmost (error ratio > +30%, red). The feedback display was shown after 300 ms of the response release and stayed for 500 ms.

**Figure 1.**
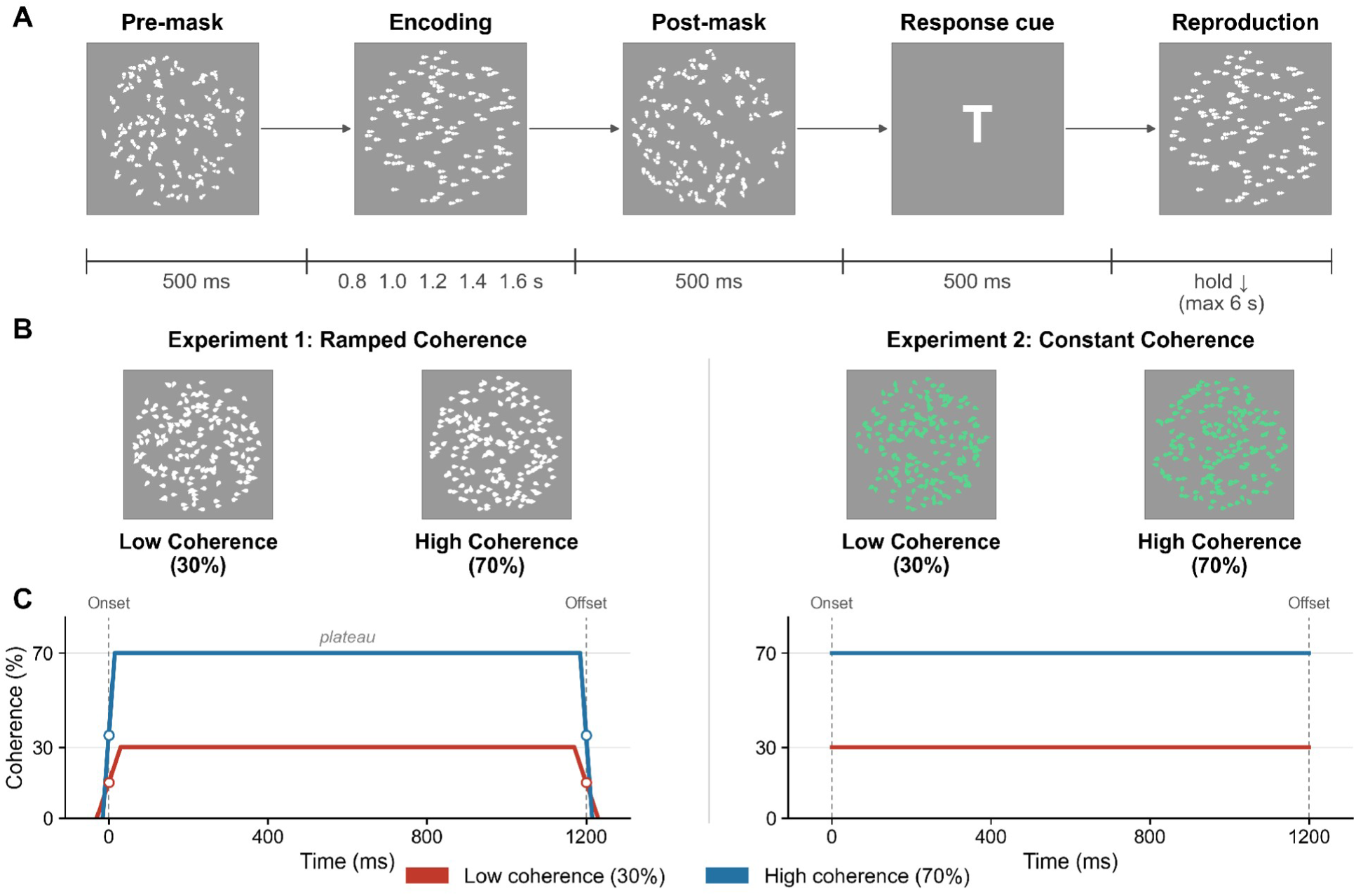
General Experimental Procedure. *Note.* (A) Trial structure: a trial starts with a pre-mask display, followed by an encoding display (30% or 70% coherence) and a post-mask. A response cue display then prompts participants to reproduce the seen duration. Dots with small arrows (not shown in the real display) represent the random-dot motion. (B) Example low- and high-coherence random-dot displays for each experiment. (C) Coherence profiles: Experiment 1 used ramped coherence (ramp-up, plateau, ramp-down), with target onset and offset defined at the midpoints of the ramp-up and ramp-down phases; Experiment 2 used constant coherence throughout each trial with sharp onsets and a distinct color cue. Dot colors in Panels B-C are for illustrative purposes.

Target durations were 0.8, 1.0, 1.2, 1.4, and 1.6 s, uniformly sampled. Motion coherence was either 30% (low coherence) or 70% (high coherence). We treat coherence as an objective manipulation of stimulus quality rather than as a direct measure of internal sensory uncertainty. The two coherence levels were randomly interleaved on a trial-by-trial basis, creating four coherence transition types that occurred with approximately equal probability (∼25% each). Repetitions and switches of coherence were thus equally likely in both experiments, and the sequence carried no information about the upcoming coherence or duration. Each participant completed 30 practice trials followed by 240 experimental trials divided into eight blocks, with short breaks between blocks.

#### 2.3.1 Experiment 1: Ramped coherence

Experiment 1 employed ramped coherence to reduce identity salience (Figure 1B-C). During encoding, motion coherence ramped up from 0% to the target level (30% or 70%), remained stable for a plateau period, and then ramped back down to 0%. Ramp durations were 60 ms for the 30% condition and 30 ms for the 70% condition; the longer ramp for the lower coherence level further increased the uncertainty of the boundary. The target interval was defined as the duration between the midpoints of the ramp-up and ramp-down phases. During reproduction, white-dot RDKs reappeared with the same motion direction. Visual feedback was provided only during practice trials.

#### 2.3.2 Experiment 2: Constant coherence

In Experiment 2, motion coherence changed immediately and remained constant throughout each trial at either 30% or 70% (Figure 1B-C). Coherence levels were randomly interleaved across trials. The encoding-phase RDK used green dots, whereas the pre-mask and post-mask phases used white dots; this color transition, together with the immediate change of the coherence level, marked each encoding phase as a distinct perceptual event. The coupled color and coherence changes were intended to make transitions more perceptually salient. During reproduction, green-dot RDKs reappeared at the same coherence level as the encoding phase.

## 3 DATA ANALYSIS

### 3.1 Preprocessing

To minimize artifacts of initial trial adaptation, the first trial of each block (8 trials total) was excluded. Outlier trials were removed if (a) reproduction errors exceeded ±3 SD within each participant × duration cell, or (b) absolute reproduction errors exceeded 0.6 s. Only participants with trial retention rates above 80% were included for formal analyses. After preprocessing, Experiment 1 participants retained ≥196 trials (*M* = 221.2, *SD* = 12.7; 92.2% retention); Experiment 2 participants retained ≥193 trials (*M* = 219.0, *SD* = 13.8; 91.2% retention).

### 3.2 Quantifying central tendency and serial dependence

Reproduction error was defined as reproduced duration minus target duration. To estimate central tendency and serial dependence jointly, while avoiding the contamination that arises when each is modeled in isolation, the primary inferential models included current and previous duration simultaneously. At the trial level, we modeled current reproduction error as

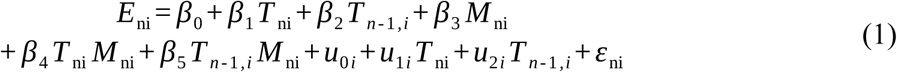

where *i* indexes participants, *T*_ni_ and *T_n_*_-1*,i*_ are the centered current and previous target durations, *u*_0*i*_ is a participant random intercept, *u*_1*i*_ and *u*_2*i*_ are participant random slopes for current and previous duration, and *M* is the moderator of interest: current low coherence in Experiment 1 and Same transition in Experiment 2. We coded *M* as a 0/1 indicator: in Experiment 1, *M* = 1 on low-coherence trials (high coherence is the reference); in Experiment 2, *M* = 1 on Same trials (Switch is the reference). The reference-condition coefficients (*β*_1_, *β*_2_) therefore describe high-coherence (Experiment 1) or Switch (Experiment 2) trials, and the interactions (*β*_4_, *β*_5_) describe the change in slope on low-coherence (Experiment 1) or Same (Experiment 2) trials. The coefficient *β*_1_ captures central tendency related to the current duration in the reference condition, *β*_2_ captures serial dependence in the reference condition, *β*_4_ tests whether the central tendency slope is modulated by the moderator, and *β*_5_ tests whether serial dependence is modulated by the moderator after controlling for current duration effects. Estimating these two interactions simultaneously allows the central tendency (current duration) and serial dependence (previous duration) channels to be modulated independently, so that any apparent moderation of serial dependence is not confounded with moderation of central tendency. Positive previous duration slopes indicate attraction toward the previous duration; negative slopes indicate repulsion.

### 3.3 Transition types

Trials were categorized by the coherence transition between consecutive trials. We label each trial’s coherence level as L (low coherence, 30%) or H (high coherence, 70%), yielding four transition types: LL (both low coherence), LH (low-to-high coherence), HL (high-to-low coherence), and HH (both high coherence); the first letter always refers to the previous trial. Same transitions comprised trials in which coherence remained constant (LL, HH); Switch transitions comprised trials in which coherence changed (LH, HL). Because coherence levels were randomly interleaved, each transition type occurred with approximately equal probability (∼25% of trials).

In these terms, the three accounts in the Introduction predict different orderings of serial dependence across the four transitions. Static reliability weighting, in which the two trials are weighted by their reliabilities, predicts HL > LL ≈ HH > LH. An account in which only current uncertainty matters predicts HL ≈ LL > HH ≈ LH. An account in which the trace is carried forward only while the perceptual context continues predicts stronger serial dependence on Same than on Switch transitions.

### 3.4 Statistical analysis

The primary tests used the trial-level linear mixed-effects model (LMM) of Equation 1, fitted to each experiment with its own moderator: current low coherence in Experiment 1 and the Same transition in Experiment 2 (Appendix A.1). The two key tests were the current duration × moderator interaction (*β*_4_), which asks whether central tendency changed with the moderator, and the previous duration × moderator interaction (*β*_5_), which asks whether serial dependence did. As a sensitivity analysis and for descriptive condition summaries, we repeated the same current-plus-previous duration regression at the participant level and compared each participant’s controlled previous-duration slopes across conditions. Effect sizes for these participant-level contrasts are reported as Cohen’s *d*.

Four further analyses complemented the primary models. First, to compare the same four transitions in both experiments, a common model let both duration slopes interact with previous coherence, current coherence, and their interaction (Appendix A.1). From it we derived the previous-duration slope of each transition, the current-coherence, previous-coherence, and Same/Switch contrasts, and two follow-up contrasts formulated after inspecting the pattern (LL against the mean of the other three transitions, and HL against LL). Second, to separate stimulus history from response history, we added the errors of the five preceding responses as predictors and let current coherence interact with the previous duration and with the previous response error (Appendix A.2). Third, we examined how coherence affected the reproductions themselves (Appendix A.3). The current Low − High contrast in mean reproduction was estimated in each experiment from the four-transition model at current and previous durations of 1.2 s, averaging equally over previous coherence. Response variability was measured as the within-participant SD of the reproductions at each duration and coherence level, averaged over durations. Fourth, the three predicted orderings (Section 3.3) were compared with Bayes factors on participant-level slopes (Appendix A.4).

## 4 RESULTS

### 4.1 Serial dependence, not central tendency, carries the main moderator effects

The primary behavioral question was whether coherence and coherence transitions changed the influence of recent history or instead reflected broader changes in duration reproduction. We therefore used dual-channel trial-level LMMs that entered current duration and previous duration simultaneously. The current-duration coefficient indexed central tendency, whereas the previous-duration coefficient indexed serial dependence after controlling for central tendency. This model allowed the moderator to act on both channels, making it possible to test whether the critical effects were specific to recent-trial carryover.

Both experiments showed the expected central tendency pattern: participants overestimated shorter durations and underestimated longer durations (Figure 2A-B). Crucially, however, the experimental moderators did not reliably change the central-tendency slope in the trial-level models. In Experiment 1, current duration strongly predicted reproduction bias on high-coherence trials, *b* = −0.386, *SE* = 0.042, *p* < .001, and on low-coherence trials, *b* = −0.371, *SE* = 0.042, *p* < .001. The current-duration × low-coherence interaction was not reliable, *b* = 0.015, *SE* = 0.020, *z* = 0.74, *p* = .460. In Experiment 2, current duration also produced strong central tendency on Switch trials, *b* = −0.487, *SE* = 0.035, *p* < .001, and Same trials, *b* = −0.486, *SE* = 0.036, *p* < .001, with no reliable current-duration × Same/Switch interaction, *b* = 0.001, *SE* = 0.022, *z* = 0.03, *p* = .977. Thus, central tendency was robust, but it was not the main channel through which the moderators acted.

**Figure 2.**
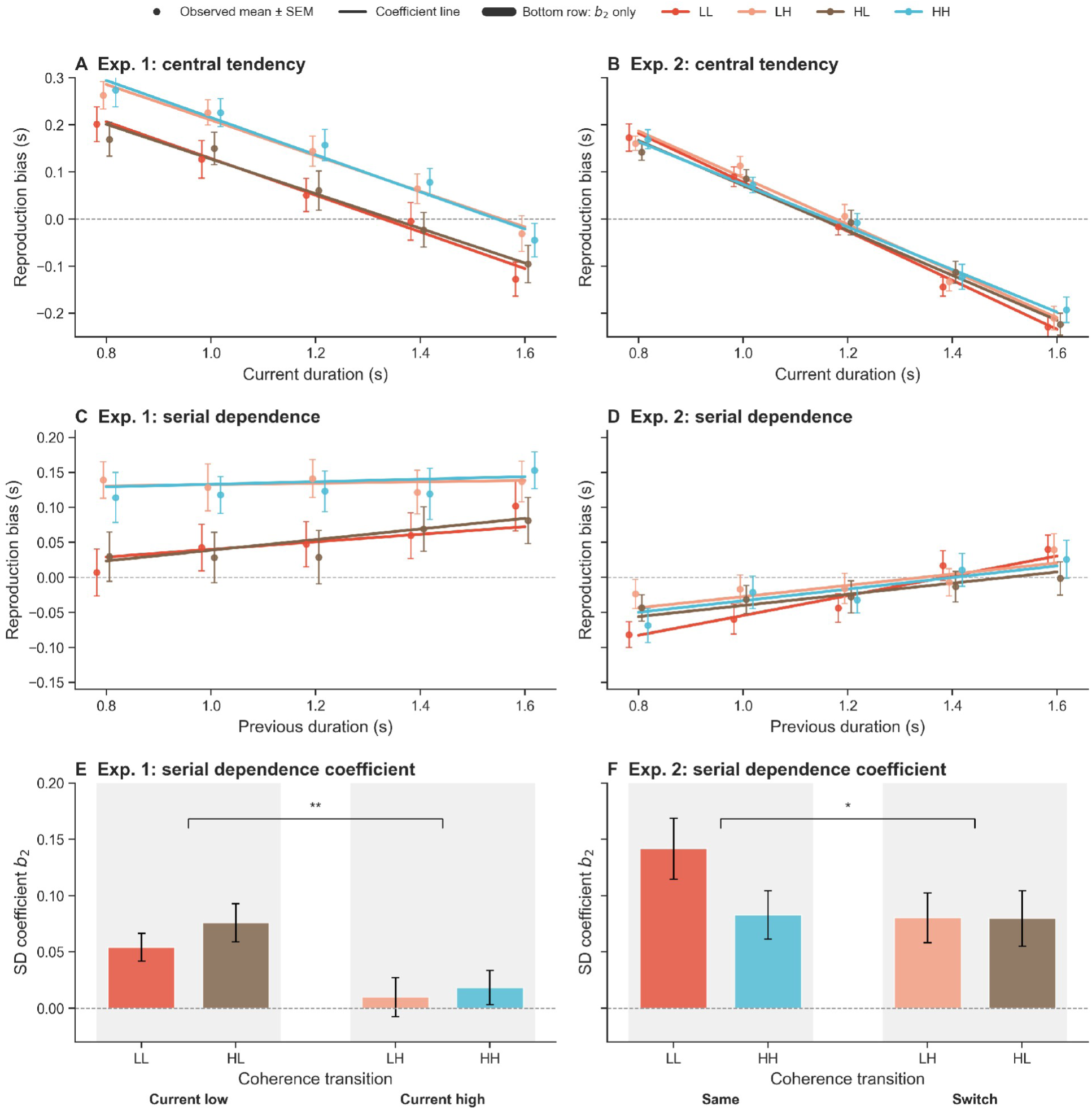
Central Tendency and Serial Dependence Across Coherence Transition Conditions. *Note.* Observed reproduction bias and coefficient-based estimates for central tendency and serial dependence. (A, B) Reproduction bias as a function of current duration in Experiments 1 and 2. Points and error bars show observed participant-averaged means ± SEM; lines show estimates reconstructed from participant-level joint regressions. (C, D) Reproduction bias as a function of previous duration, using the same error-bar convention: participant-averaged means ± SEM. (E) Experiment 1 controlled serial-dependence coefficients (*b*_2_) grouped by current coherence. Bars and error bars show participant-level coefficient means ± SEM. Current-low trials comprise LL and HL transitions; current-high trials comprise LH and HH transitions. (F) Experiment 2 controlled serial-dependence coefficients grouped by Same versus Switch transitions, again with participant-level means ± SEM. Same trials comprise LL and HH transitions; Switch trials comprise LH and HL transitions. Stars in panels E and F mark participant-level planned contrasts on controlled *b*_2_ estimates (\**p* < .05, \*\**p* < .01). L denotes low coherence (30% coherence), and H denotes high coherence (70% coherence).

In contrast, Experiment 1 showed a selective coherence modulation of serial dependence (Figure 2C,E). The previous-duration slope was small and not reliable on high-coherence trials, *b* = 0.019, *SE* = 0.014, *p* = .181, but was larger on low-coherence trials, *b* = 0.066, *SE* = 0.014, *p* < .001. In the primary trial-level mixed model, the previous-duration × low-coherence interaction was reliable, *b* = 0.048, *SE* = 0.020, *z* = 2.37, *p* = .018. As a robustness check, we repeated the same current-plus-previous duration regression at the participant level and compared each participant’s controlled previous-duration slopes across coherence conditions. This produced the same effect-size pattern: slopes were larger on current-low than current-high trials, mean difference = 0.0477, *t*(21) = 3.52, *p* = .002, *d* = 0.751. Lower current coherence therefore increased the influence of recent temporal history while leaving the central-tendency channel essentially unchanged.

The four-transition model showed the same pattern at each level of previous coherence (Appendix A.1). The previous-duration slopes were HL = 0.077 and LL = 0.054 on current-low trials, against HH = 0.028 and LH = 0.009 on current-high trials. Averaged over previous coherence, the current Low − High difference was 0.047, 95% CI [0.008, 0.086], *p* = .020, whereas neither previous coherence (High − Low = 0.021, *p* = .298) nor Same versus Switch (−0.002, *p* = .931) had a reliable effect. In the comparison of the predicted orderings, grouping by current coherence was favored over static reliability weighting (Bayes factor = 6.4; Appendix A.4).

Experiment 2 showed a different pattern (Figure 2D,F). Previous duration reliably predicted current reproduction bias on Switch trials, *b* = 0.079, *SE* = 0.019, *p* < .001, and the estimated slope was numerically larger on Same trials, *b* = 0.108, *SE* = 0.019, *p* < .001. The planned trial-level previous-duration × Same/Switch interaction did not reach significance, *b* = 0.029, *SE* = 0.022, *z* = 1.33, *p* = .182, although the participant-level controlled-slope contrast was positive, mean difference = 0.0279, *t*(21) = 2.26, *p* = .034, *d* = 0.483. The four-transition model shows why the two-category contrast was weak: the slope was 0.136 on LL trials, but 0.078, 0.079, and 0.079 on HL, LH, and HH trials. The elevation was confined to repeated low coherence, which exceeded the mean of the other three transitions by 0.057, 95% CI [0.008, 0.107], *p* = .024 (Bonferroni-adjusted over the four follow-up contrasts, *p* = .096). Neither the current-coherence contrast (0.028, *p* = .203) nor the previous-coherence contrast (−0.029, *p* = .176) was reliable. In the comparison of orderings, an elevation on LL alone was favored over grouping by current coherence (Bayes factor = 60.8). Static reliability weighting, which predicts the strongest dependence on HL trials, was not the favored ordering in either experiment.

A complementary analysis examined response history, because the reproduction display resembled the stimulus and each response may itself leave a trace (Appendix A.2). The errors of the five preceding responses (reproduced minus physical duration) were entered together with current and previous physical duration. Previous response errors impacted the current error across several lags in both experiments, indicating a slowly changing response bias. The coherence modulation of serial dependence in Experiment 1 remained when these response-history terms were included, previous duration × coherence *b* = 0.088, *SE* = 0.028, *p* = .002, whereas the interaction of the previous response error with coherence was not reliable (*b* = 0.052, *p* = .093). The coherence effect therefore concerns the history of the stimuli beyond the history of the responses; the computational model below represents the two sources separately.

Coherence also affected the reproductions themselves, apart from their dependence on history. In Experiment 1, reproductions were shorter on current-low than on current-high trials by 84 ms (−0.084 s, 95% CI [−0.095, −0.073], *p* < .001; Figure 2A). In Experiment 2 this shift was small (−0.012 s, 95% CI [−0.024, 0.000], *p* = .052), and the two experiments differed significantly (difference = −0.073 s, *p* < .001). The shift in Experiment 1 was additive: it was about the same at all five durations (−83, −87, −88, −84, and −72 ms from 0.8 to 1.6 s) and did not scale with duration (Appendix A.3). Reproductions were also more variable under low coherence in Experiment 1: the within-participant SD at a given duration was 0.193 s on low-and 0.173 s on high-coherence trials, difference = 0.021 s, 95% CI [0.011, 0.031], *t*(21) = 4.24, *p* < .001. In Experiment 2 the difference was small and not reliable (0.206 vs. 0.201 s; difference = 0.005 s, *t*(21) = 1.49, *p* = .152). Coherence thus changed the variability of the reproductions in Experiment 1, but not in Experiment 2 (Appendix A.3).

### 4.2 Cross-experiment comparison of coherence and transition effects

The two experiments showed related but not identical moderator profiles (Figure 3). We compared participant-level controlled previous-duration slopes by Same/Switch status and by current coherence to visualize whether the two designs produced convergent modulation. These analyses were descriptive because the experiments were run on separate samples and differed in coherence dynamics and perceptual salience.

**Figure 3.**
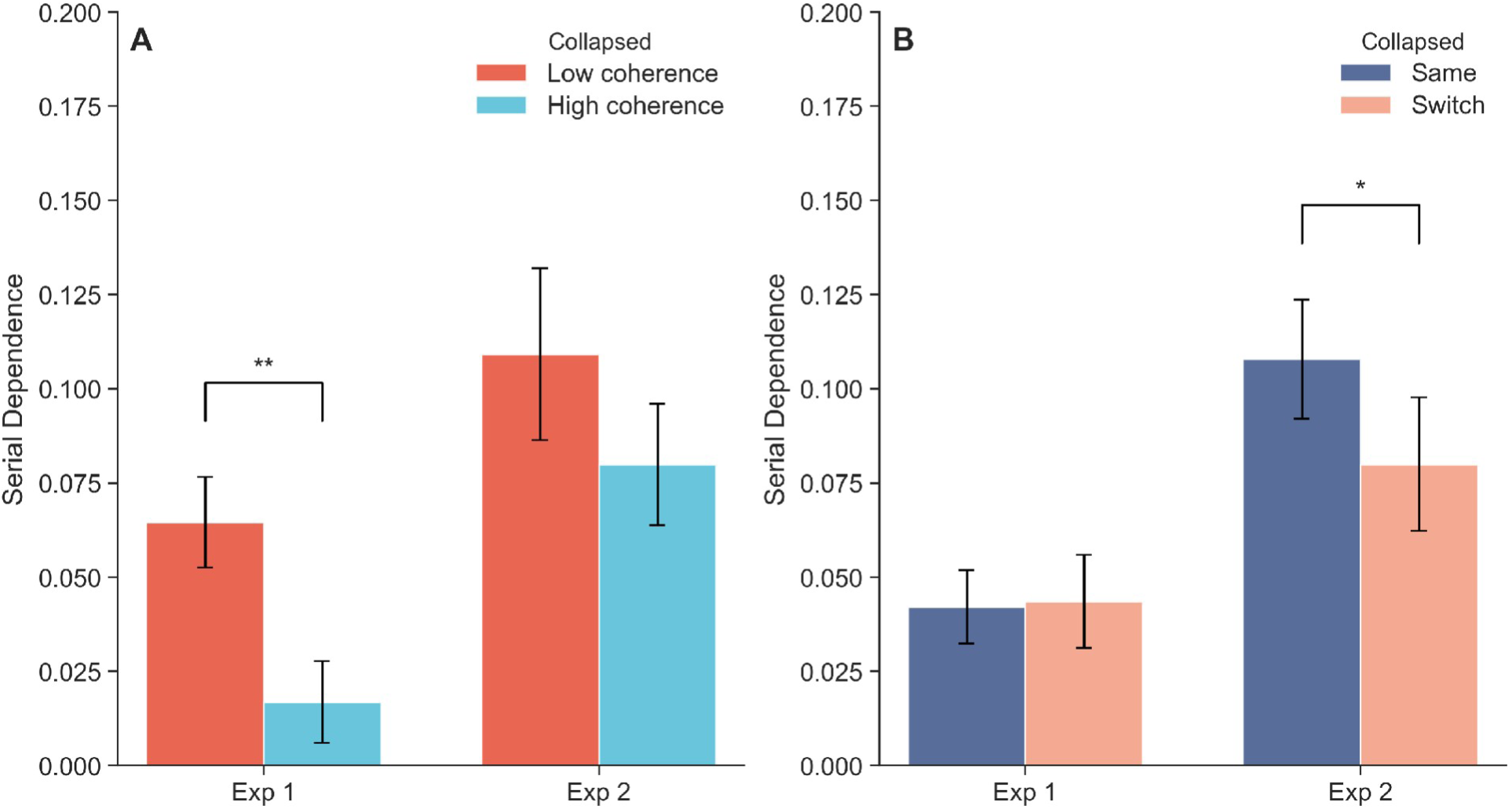
Cross-Experiment Comparison of Controlled Serial Dependence Slopes. *Note.* Controlled previous-duration slopes compared across experiments after collapsing the four coherence transitions into the contrast of interest. (A) Collapsed current-coherence contrast: Low versus High current coherence. (B) Collapsed transition-consistency contrast: Same versus Switch transitions. Error bars represent ± SEM. Stars denote participant-level paired comparisons of the controlled slopes shown in each panel (\**p* < .05, \*\**p* < .01); these comparisons are distinct from the primary LMM tests.

For current coherence (Figure 3A), Experiment 1 showed stronger slopes on low- than high-coherence trials, *t*(21) = 3.52, *p* = .002, whereas Experiment 2 did not, *t*(21) = 1.24, *p* = .230. For Same/Switch status (Figure 3B), Experiment 1 showed no reliable difference, *t*(21) = −0.10, *p* = .919, whereas Experiment 2 showed stronger controlled previous-duration slopes on Same than Switch trials, *t*(21) = 2.26, *p* = .034. Current evidence quality thus produced the clearest modulation of serial dependence in Experiment 1, whereas the repetition of coherence produced a weaker pattern in Experiment 2, which the four-transition analysis locates on LL trials.

### 4.3 Temporal locality of the carryover effect

As a robustness check, we asked whether the serial-dependence effects were temporally local. We regressed current reproduction errors against durations from trials n−1 through n−3, together with forward-looking checks at n+1 and n+2 (Figure 4). This analysis tested whether the behavioral effects were concentrated in recent history rather than reflecting broad temporal drift.

**Figure 4.**
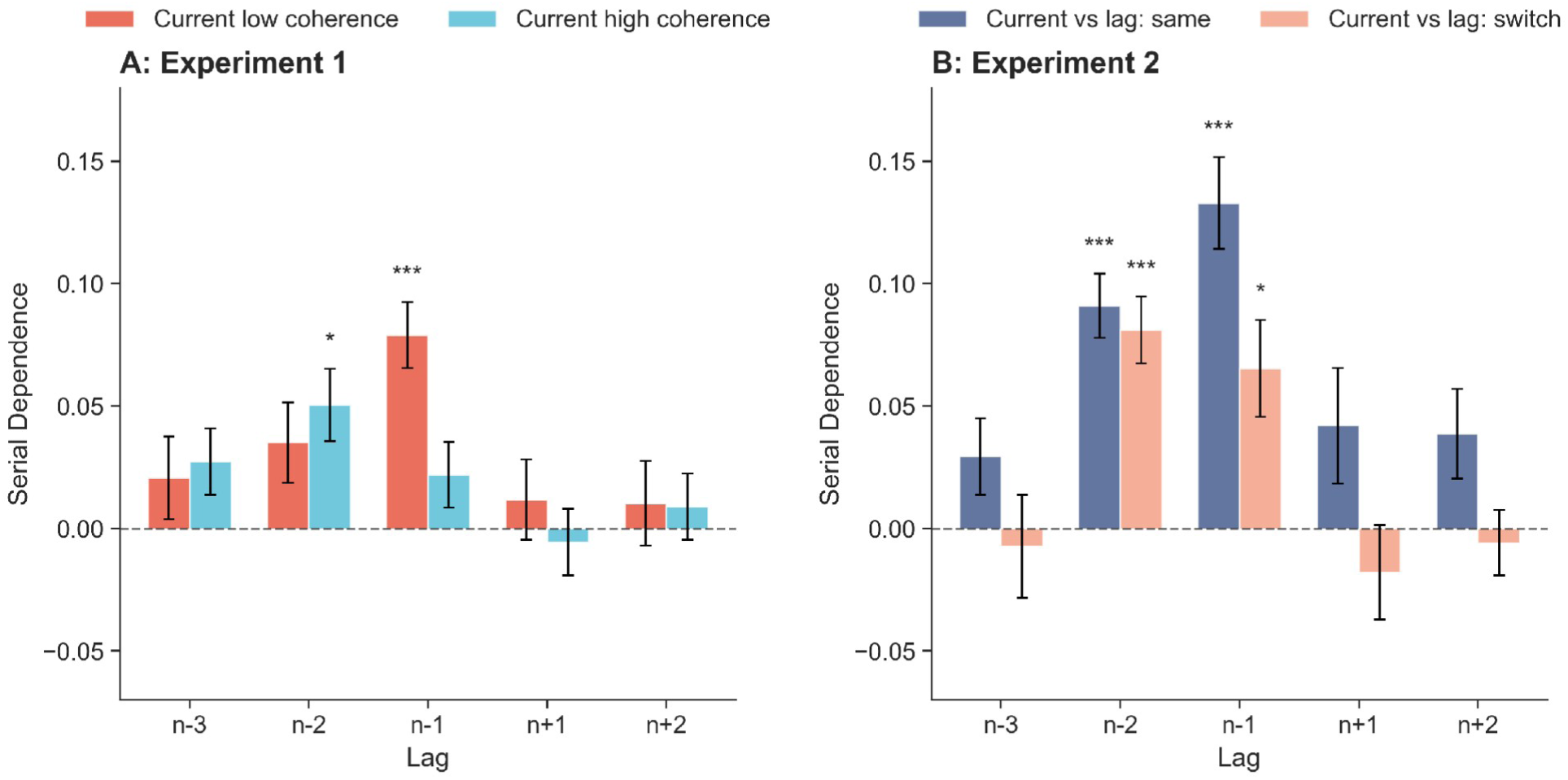
Temporal Window of Serial Dependence. *Note.* Serial dependence coefficients (*b*) for preceding trials (n−1, n−2, n−3) and future trials (n+1, n+2). (A) Experiment 1, grouped by current coherence (Low vs. High). (B) Experiment 2, grouped by lag-specific transition status: Same means the current trial and the trial shown on the x-axis had the same coherence/context; Switch means they differed. Thus the n−2 bars compare current vs. n−2, not current vs. n−1. Error bars represent ± SEM. Significance markers use Bonferroni-corrected *p* values across the five lag tests within each condition: \*\*\**p* < .001, \*\**p* < .01, \**p* < .05.

In Experiment 1, low-coherence trials showed reliable attraction only at n−1, *b* = 0.079 ± 0.013, *t*(21) = 5.89, Bonferroni-corrected *p* < .001; all other lags were nonsignificant after correction, corrected *p*s ≥ .221. High-coherence trials showed a reliable effect at n−2, *b* = 0.050 ± 0.015, *t*(21) = 3.41, corrected *p* = .013, whereas the n−1 effect was not reliable after correction, *b* = 0.022 ± 0.013, *t*(21) = 1.64, corrected *p* = .576; all other lags were nonsignificant, corrected *p*s ≥ .283.

In Experiment 2, Same/Switch status was recomputed separately for each lag, so each bar compares the current trial with the specific lagged or future trial shown on the x-axis. Lag-specific Same trials showed reliable attraction at n−1, *b* = 0.133 ± 0.019, *t*(21) = 7.08, Bonferroni-corrected *p* < .001, and n−2, *b* = 0.091 ± 0.013, *t*(21) = 6.94, corrected *p* < .001. Lag-specific Switch trials also showed attraction at n−1, *b* = 0.065 ± 0.020, *t*(21) = 3.29, corrected *p* = .018, and n−2, *b* = 0.081 ± 0.014, *t*(21) = 5.94, corrected *p* < .001. Effects at n−3 and the future-trial checks were not reliable after correction for either Same or Switch trials, corrected *p*s ≥ .237 for Same and ≥ .999 for Switch. Serial dependence was therefore concentrated at the recent lags, with no corresponding future-trial pattern.

## 5 COMPUTATIONAL MODELING

The behavioral results pose three constraints for a model. Current coherence changed serial dependence but not central tendency (Experiment 1); serial dependence was elevated when low coherence repeated (Experiment 2); and responses carried a slow bias across several trials. We asked whether one sequential observer can produce all three, using a three-state Kalman filter based on the iterative updating framework of Glasauer and Shi (2022). A Kalman filter describes how an observer combines a current observation with prior internal states, weighting each source by its expected reliability. This framework has been used to explain both central tendency and serial dependence in magnitude perception (Glasauer & Shi, 2022; Petzschner et al., 2015; Shi et al., 2013).

### 5.1 Model architecture

The model assumes that observers carry three internal states from one trial to the next (Figure 5; Appendix B.1). The *fast state* is the current duration estimate. The *slow state* tracks the mean of the durations encountered so far and therefore produces central tendency. The *bias state* is a slowly drifting response bias.

**Figure 5.**
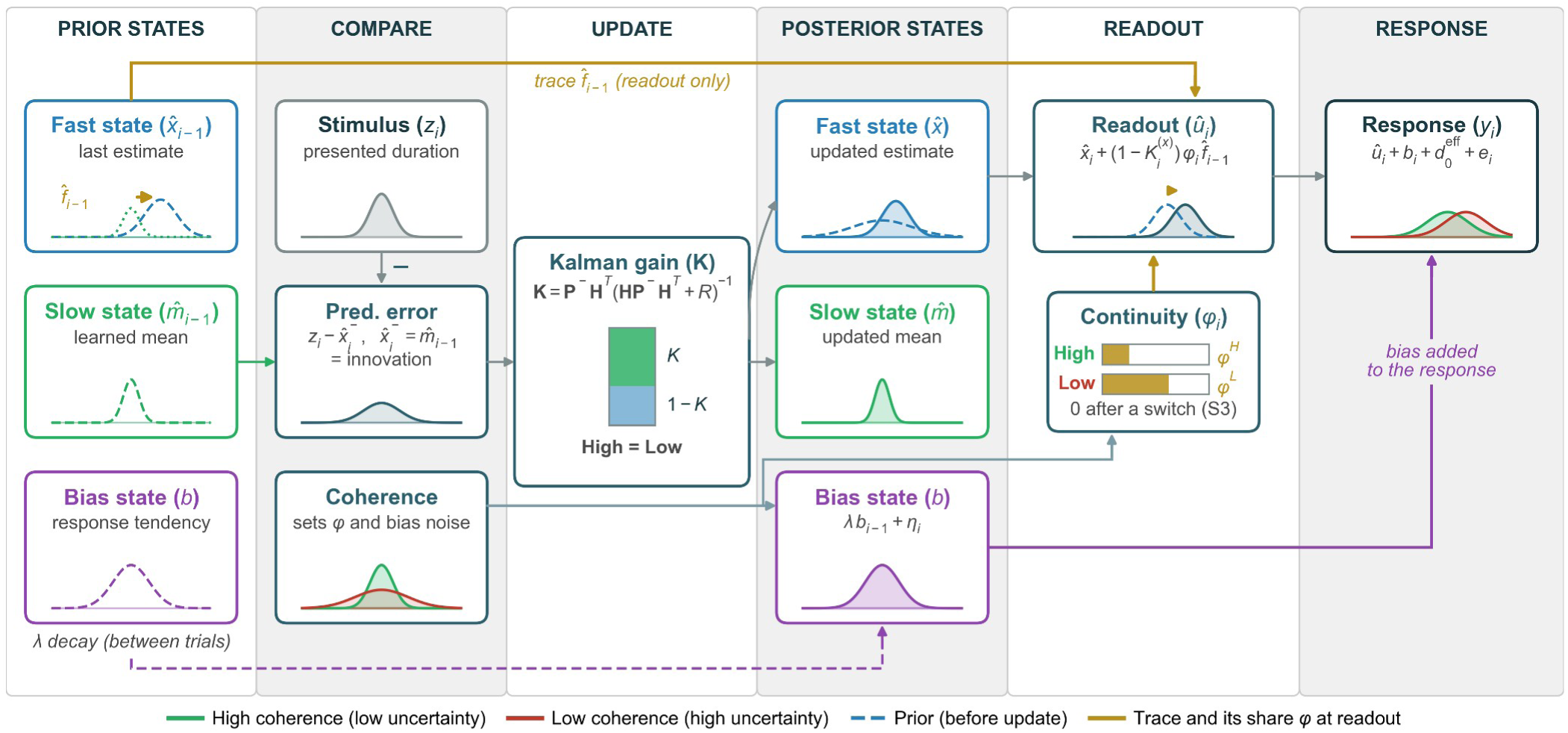
Three-State Kalman Filter Model. *Note.* Schematic of the three-state Kalman filter on a single trial. The observer predicts the current duration from the learned mean (*m̂*, slow state) and combines the prediction with the presented duration (*s_i_*) according to the Kalman gain (*K*^(^*^x^*^)^), which gives the duration estimate (*x̂*, fast state). At readout, a share *φ* of the fast deviation of the previous trial (*f̂_i_* _- 1_) is added to the prediction; *φ* depends on the current coherence and can be set to zero when coherence switches. The response adds a slowly drifting bias (*b*, decaying at rate *λ*), a coherence-dependent offset, and noise. The figure depicts the selected C3-B2 specification, in which coherence does not change the filter gain and the response bias evolves outside the duration filter. The bar lengths and response distributions are illustrative; the model does not impose an ordering of *φ^L^* and *φ^H^* .

On each trial the observer combines a prior with the presented duration *s_i_*. In the filter itself, the prior is the slow state *m̂_i_* _- 1_, the learned mean of the stimulus distribution, and the estimate is 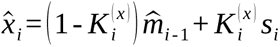. The Kalman gain 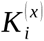 is the weight of the presented duration and 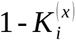 the weight left for the prior, so a larger gain means that the estimate follows the current duration more closely. The learned mean then moves toward the presented duration with a smaller gain, 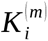. These two states and their updates follow the two-state model of Glasauer and Shi (2022), in which the previous duration influences the current estimate only through the learned mean.

The present model adds a second route, the trace of the previous trial itself. We take this trace to be the previous duration estimate, *x̂_i_* _- 1_. The estimate that is reproduced, which we call the readout *û_i_*, is formed from a prior for the current trial that lies between the learned mean and the trace, (1 - *φ_i_*) *m̂_i_* _- 1_ + *φ_i_ x̂_i_* _- 1_. The retention *φ_i_* is the weight of the trace in this prior. Combining the prior with the presented duration at the gain of the filter gives

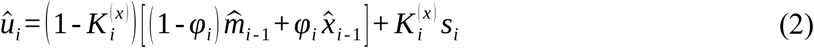

With *φ_i_* = 0 the prior is the learned mean alone and the readout is the estimate of the two-state model, *x̂_i_*. With *φ_i_* = 1 the prior is the previous estimate. Equation 2 can also be written as 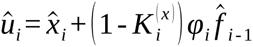, where *f̂_i_*_-1_ = *x̂_i_*_-1_- *m̂_i_*_-1_ is the deviation of the previous estimate from the learned mean (the fast deviation in Figure 5 and Appendix B.1).

The prior as a whole receives the weight 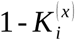, so the readout remains precision weighted. The retention can differ between low and high current coherence (*φ^L^*, *φ^H^*), because the readout is formed after the stimulus has been presented and the current coherence is known at that point. The readout is not written back into the duration filter: *x̂_i_* and *m̂_i_* are updated without it, while the response-bias state evolves separately, so the trace does not accumulate over trials. The response is the readout plus an offset, which may differ between coherence levels to capture the mean shift reported in Section 4.1, plus the response bias and response noise.

This architecture separates two ways in which coherence could act. If low coherence lowers the gain *K*^(^*^x^*^)^, it strengthens central tendency and serial dependence together, because both scale with 1 - *K*^(*x*)^. If low coherence raises the retention *φ*, it changes serial dependence only: the slope of the response on the current duration is *K*^(^*^x^*^)^ whatever the value of *φ* (Appendix B.2).

### 5.2 Model space and comparison

We compared 148 candidate models that answer three questions (Appendix B gives the full technical specification). First, where does coherence change the observer’s noise (C)? Coherence could modulate the process noise of the fast state (*q*_1_), of the slow state (*q*_2_), or of the bias state (*q*_3_), the measurement noise (*R*), or any non-empty combination of these components (15 levels). Process noise is the amount by which the observer expects a state to change from one trial to the next; measurement noise is the unreliability that the observer assigns to the current duration. A modulation of *q*_1_ or *R* changes the gain on the current trial. Second, what happens when coherence switches between trials (S)? The model could include no switch effect (S0), a partial reset of the slow state toward the overall mean (S1), a reduced Kalman gain (S2), or a reset of the trace, so that the previous deviation is not retained on switch trials (*φ* = 0; S3). Third, where does the slow bias act (B)? There could be no bias state (B0), a bias that enters the observer’s prediction (B1), or a bias that is added to the response without being tracked by the observer (B2). Every model had *φ^L^* and *φ^H^* as free parameters. Each level was thus tested on top of the retention mechanism, and a model that does not need retention can set both to zero.

Models were fitted separately for each participant by maximum likelihood and compared using the Akaike Information Criterion (AIC), averaged over participants. AIC measures the fit to all responses and is not specific to the effects of interest. Each model was therefore also simulated on every participant’s own trial sequence and assessed on whether it reproduced the focal contrast of its experiment (current Low − High in Experiment 1; LL against the other three transitions in Experiment 2), central tendency, and response variability.

The three dimensions gave answers of different strength (Table 1). The bias dimension was decisive: models with a response-level bias (B2) were better than all models with a prediction-level bias or without a bias state by at least 21 AIC units in Experiment 1 and 15 in Experiment 2. On the switch dimension, Experiment 1 favored no switch effect (S0), and no model with the trace reset reproduced its Low − High contrast. In Experiment 2, the S0 models had the lowest AIC, but none of them reproduced the elevation on LL trials; all 37 models with the trace reset (S3) did, against 2 of the 111 other models, at a cost of 0.56 AIC units for the best S3 model. The coherence dimension was not identified. The best models of several C levels lay within a fraction of an AIC unit of each other, and no C level produced the effect of current coherence: at the fitted parameters, the weight left for the prediction, 1 - *K*^(^*^x^*^)^, differed between low and high coherence by at most 3% in Experiment 1 and 6% in Experiment 2 (Appendix B.3). We therefore report C3-S0-B2 for Experiment 1 and C3-S3-B2 for Experiment 2. C3, in which coherence modulates only the drift of the response bias, is among the levels with the fewest parameters, lies within 0.12 and 0.56 AIC units of the best model, and reproduces response variability; in these two models, coherence does not enter the observer’s filter at all.

**Table 1.** Model Comparison by Dimension.

| Dimension | Level | Models | Exp. 1: best ΔAIC | Exp. 1: reproduce focal contrast | Exp. 2: best ΔAIC | Exp. 2: reproduce focal contrast |
|---|---|---|---|---|---|---|
| Bias | B0, no bias state | 28 | 21.0 | 12 | 15.1 | 7 |
|  | B1, in the prediction | 60 | 23.6 | 19 | 16.4 | 17 |
|  | B2, in the response | 60 | 0.0 | 33 | 0.0 | 15 |
| Switch | S0, no switch effect | 37 | 0.0 | 21 | 0.0 | 0 |
|  | S1, reset of the mean | 37 | 1.4 | 23 | 1.3 | 2 |
|  | S2, reduced gain | 37 | 1.8 | 20 | 1.5 | 0 |
|  | S3, reset of the trace | 37 | 0.6 | 0 | 0.6 | 37 |

*Note.* ΔAIC is the mean AIC per participant relative to the best of the 148 models of the experiment; the table gives the best model of each level. The last column of each experiment counts the models of the level whose simulations reproduced the focal contrast (Experiment 1: current Low − High serial dependence; Experiment 2: LL against the other three transitions) within 1.5 standard errors of the data.

### 5.3 Parameter estimates and the origin of serial dependence

The fitted parameters describe how the selected models generate the behavioral patterns (Figure 6). The gain on the current duration did not differ between low and high coherence (Experiment 1: 0.61 and 0.61; Experiment 2: 0.51 and 0.51), and neither did the gain on the mean (0.11 and 0.11; 0.20 and 0.20; all *p* > .38; Figure 6A,D). The models therefore predict the same central tendency under both coherence levels, as observed. Experiment 2 had the lower gain on the current duration and the higher gain on the mean, which gives its stronger central tendency and its larger overall serial dependence.

**Figure 6.**
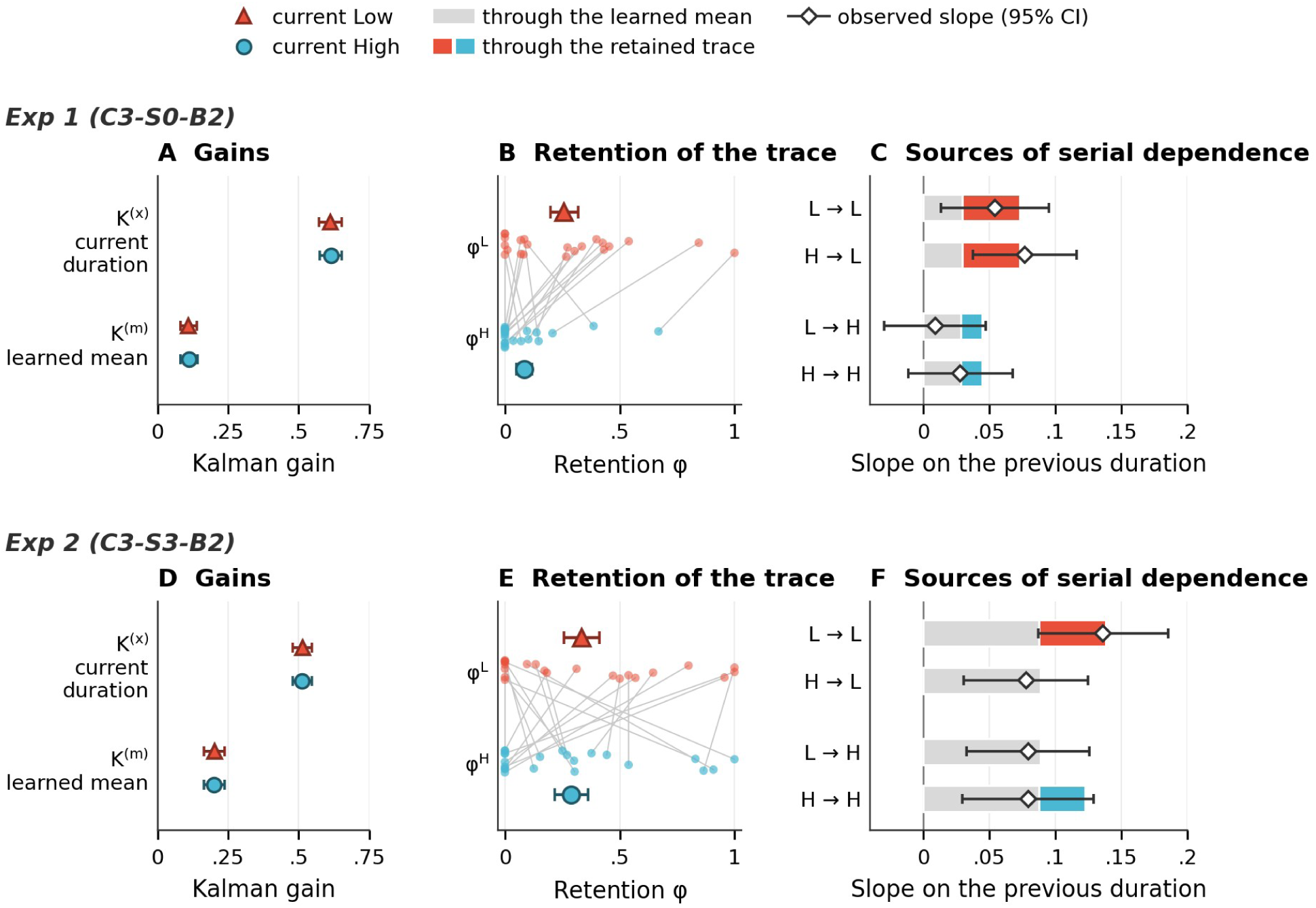
Gains, Retention, and Sources of Serial Dependence in the Selected Models. *Note.* Top row: Experiment 1 (C3-S0-B2). Bottom row: Experiment 2 (C3-S3-B2). Red triangles denote low and blue circles high current coherence. (A, D) Kalman gain on the current duration (*_K_*^(^*^x^*^)^) and on the learned mean (*_K_*^(^*^m^*^)^); participant means ± SEM. (B, E) Retention of the trace under low and high current coherence (*φ^L^*, *φ^H^*). Small dots are participants, with a line joining the two estimates of each participant; large symbols are participant means ± SEM. (C, F) Model slope of the response on the previous duration for each transition (previous → current coherence), divided into the route through the learned mean (gray) and the route through the retained trace (colored by current coherence). Slopes are computed at the fitted parameters and averaged over the trials of each transition and over participants. Diamonds show the observed slopes of the four-transition mixed model with 95% confidence intervals (Appendix A.1). The remaining parameter estimates are given in Appendix B.3 (Table B2).

Coherence acted on retention. In Experiment 1, the retained share of the previous trace was larger on low- than on high-coherence trials, *φ^L^* = 0.26 against *φ^H^* = 0.08 (medians 0.18 and 0; larger in 14 of 22 participants; Wilcoxon signed-rank *p* = .003; Figure 6B,E). In Experiment 2 the two did not differ (0.34 and 0.29, *p* = .70); here retention depended on the transition, through the reset at a switch of coherence.

To see where serial dependence comes from, we separated the model’s slope on the previous duration into the route through the learned mean and the route through the retained trace (Figure 6C,F; Appendix B.2). The route through the mean was the same for all four transitions (0.029 in Experiment 1 and 0.088 in Experiment 2). In Experiment 1, the retained trace added 0.044 when the current trial was low coherence and 0.016 when it was high, whatever the previous coherence. In Experiment 2, the retained trace added 0.050 on LL and 0.035 on HH trials, and nothing on the two switch transitions.

The bias state decayed slowly (*λ* = 0.72 and 0.78) and drifted by a few tens of milliseconds per trial (Appendix B.3, Table B2). It accounts for the dependence of the responses on the errors of several preceding responses (Section 4.1) and does not contribute to the slope on the previous duration.

Recovery simulations distinguished S0 from S3 at the cohort level under the fitted generating conditions, whereas individual retention values, especially under high coherence, were weakly identified. Appendix B.5–B.6 reports parameter and coherence-effect recovery, the pairwise model-recovery comparison, and profile-likelihood diagnostics.

### 5.4 Predictive checks

To assess whether the models reproduced the key behavioral phenomena, we simulated responses from the fitted parameters. In each of 100 simulated cohorts, every participant was simulated once on their own trial sequence, and the simulated responses were analyzed like the data. Figure 7A–D shows reproduction bias as a function of the previous duration, and Figure 7E–G summarizes central tendency, serial dependence, and response variability by current coherence. Response variability is the residual SD of the regression on current and previous duration.

**Figure 7.**
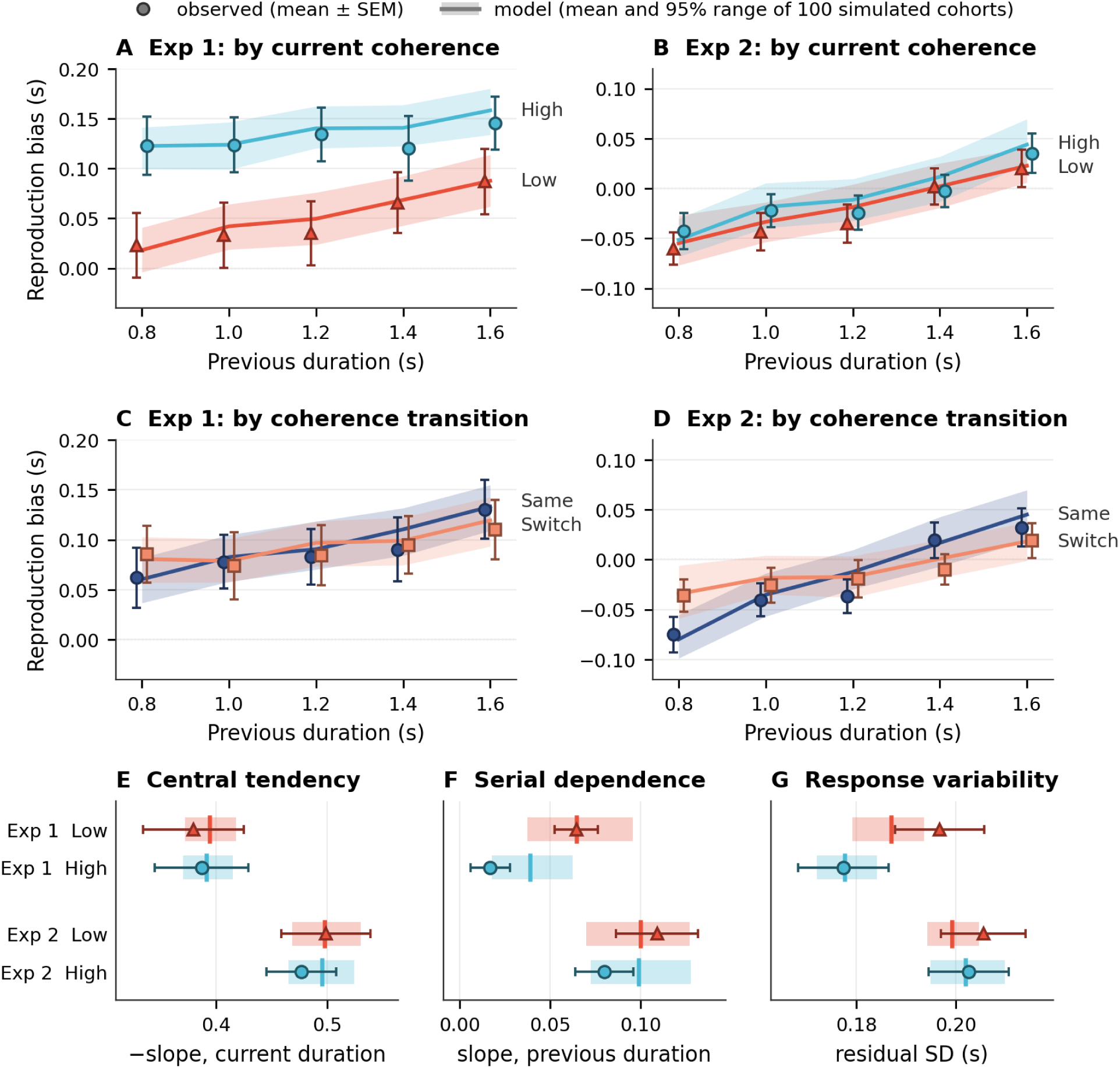
Predictive Checks: Observed Data and Model Predictions. *Note.* Symbols with error bars show observed participant means ± SEM. Lines (A–D) and vertical ticks (E–G) show the model mean over 100 simulated cohorts, and shaded regions the central 95% range across cohorts. Experiment 1 is simulated with C3-S0-B2 and Experiment 2 with C3-S3-B2. (A–D) Mean reproduction bias as a function of the previous duration, by current coherence (A, B) and by Same versus Switch transition (C, D). The vertical axes of the two experiments have the same span (0.24 s), so that slopes can be compared. (E–G) Central tendency (the negative slope on the current duration), serial dependence (the slope on the previous duration), and response variability (the residual SD of the same regression), by experiment and current coherence.

Central tendency was reproduced in both experiments (Figure 7E). Response variability was reproduced under high coherence in both experiments and under low coherence in Experiment 2; in Experiment 1 the model placed the low-coherence residual SD slightly too low (Figure 7G; 0.187, 95% range of the simulated cohorts [0.179, 0.194], against an observed 0.197 ± 0.009). Resolved by duration, the model’s residual SD was flat, whereas in Experiment 1 the observed SD grew with duration (Appendix B.4); the model has constant response noise.

For serial dependence, the Experiment 1 model reproduced the slope on low-coherence trials (0.065 against an observed 0.065) and the separation of the two coherence levels (Figure 7A), but it overestimated the slope on high-coherence trials (0.039 [0.018, 0.063] against 0.017 ± 0.011; Figure 7F). The Experiment 2 model reproduced the steeper dependence on Same than on Switch transitions (Figure 7D), which arises from the reset of the trace. It predicted the same slope for low and high current coherence (0.100 and 0.099, against an observed 0.109 and 0.080) and thus an elevation on HH trials almost as large as on LL trials (Figure 6F), which the data do not show. The remaining misfit is the same in both experiments: retention on current-high trials is overestimated.

### 5.5 A test with manipulated sequence volatility

In the present experiments the durations were drawn independently on every trial, so the volatility of the sequence did not vary. To test whether the model also accounts for a change of volatility, we applied it to the data of Glasauer and Shi (2022), in which 14 participants reproduced the same set of durations in two orders: fully randomized, and as a random walk in which successive durations were similar (Appendix C). In the randomized order, central tendency was strong and the dependence on the previous duration positive (0.455 and 0.109). In the random-walk order, central tendency nearly vanished and the dependence on the previous duration became negative (0.096 and −0.086). We fitted the Experiment 1 model to the randomized condition only and predicted the random-walk condition with all parameters fixed. The predicted central tendency and serial dependence (0.091 and −0.084) matched the data (paired *p* = .90 and .96). The effect of sequence volatility on both biases thus follows from the order of the stimuli, with no change in the observer. Appendix C compares these predictions with those of the two-state model of the original study.

## 6 GENERAL DISCUSSION

Across two experiments, the attraction of duration reproduction toward the preceding duration depended on the coherence of the current stimulus and on whether coherence repeated across trials. In Experiment 1, low current coherence strengthened serial dependence while leaving central tendency unchanged, and the coherence of the preceding trial had no reliable effect. In Experiment 2, in which each encoding interval was sharply marked, serial dependence was present on all transitions and elevated when low coherence repeated. A three-state Kalman filter accounted for these patterns with one mechanism: the observer retains part of the trace of the preceding trial in the prior for the current estimate; current coherence sets how much is retained (Experiment 1), and a change of coherence interrupts the retention (Experiment 2). The weight of the current duration itself, the Kalman gain, did not depend on coherence. More broadly, the findings speak to a general problem for adaptive cognition: how much recent experience should remain active when current evidence is weak or when the perceptual context changes?

### 6.1 Precision weighting and the retention of recent history

The usual answer to the question of how much weight the past should receive is precision weighting: the less reliable the current evidence, the more the observer relies on prior information (Ernst & Banks, 2002; Feldman & Friston, 2010; van Bergen & Jehee, 2019). In its most direct form, lower stimulus quality lowers the weight of the current evidence, that is, the gain. This form accounts for the growth of central tendency with measurement noise (Petzschner et al., 2015; Shi et al., 2013) and for the stronger serial dependence observed with uncertain stimuli (Ceylan et al., 2021; Cicchini et al., 2018; Gallagher & Benton, 2022). It also makes a joint prediction. A lower gain leaves more weight for everything the observer brings to the trial, so it should strengthen central tendency and serial dependence together. The present data did not bear this prediction out. In both experiments central tendency was the same under low and high coherence, and in the fitted models no modulation of process or measurement noise by coherence produced a difference in gain. A possible reason is that motion coherence is a property of the carrier of the interval and not of the interval itself. The onset and offset that define the duration were visible at both coherence levels, so coherence may have changed the measured duration little. By contrast, duration reproduction under memory pressure shows stronger central tendency (Zang et al., 2026), as expected when the duration measurement itself becomes less reliable.

What coherence changed was the use of the preceding trial. In the model, the prior for the current trial lies between the learned mean and the trace of the preceding trial, and coherence regulates the weight of the trace in this prior, the retention. The prior as a whole is then combined with the current duration at the same gain, so the updating remains precision weighted throughout. Coherence thus determined what the prior contains and left unchanged how the prior is weighed against the evidence. One reading is that weak current evidence leads the observer to rely on a richer prior, one that includes the most recent event in addition to the long-run mean. Experiment 1 showed this without sharp perceptual boundaries between events. Experiment 2 showed the complementary case: with sharply marked encoding events, retention did not differ between coherence levels, but it was interrupted when the coherence of the event changed (Section 6.3). This conclusion is bounded by the model space we tested. Within the 148 candidate models, the effects of coherence level and of coherence transitions were captured by one precision-weighted observer in which coherence regulates the retention of recent history.

The two biases are therefore regulated separately in this account. Central tendency and serial dependence are signatures of how the observer uses the past over a long and a short timescale (Glasauer & Shi, 2022): the gain affects both, whereas retention changes serial dependence alone. A manipulation that changes serial dependence and leaves central tendency unchanged thus points to retention, and one that changes both points to the gain.

### 6.2 Relation to context effects and the reliability asymmetry

Serial dependence is known to depend on context. Bae and Luck (2020) demonstrated that serial dependence in motion direction occurred only when motion was the reported feature, and C. Fischer et al. (2020) showed that contextual features such as serial position modulate carryover. Task relevance also determines whether attractive or repulsive sequential biases emerge (Ceylan & Pascucci, 2023), and working memory and task demands modulate opposing serial biases (Cheng, Chen, & Shi, 2024). These results have been taken as evidence that serial dependence requires task relevance or contextual continuity, in line with accounts that tie it to the continuity of objects and events (Manassi & Whitney, 2024; Pascucci et al., 2023) and to spatiotemporal routines (Pascucci & Kristjánsson, 2026). The present model offers one way to express such a requirement within an updating framework: a preceding event contributes beyond the learned mean only if its trace is retained, and retention can be conditional on continuity, as at the change of coherence in Experiment 2. Not every context effect need act in this way. Some may reflect changes in the effective precision of stimulus representations, as attention and cognitive load are known to produce, and these act through the gain. By the criterion above, they should change central tendency as well. The two kinds of effect can therefore be told apart when precision and task context are manipulated independently within one design.

The separation of gain and retention also bears on the asymmetry identified by Gallagher and Benton (2022): serial dependence is modulated by reliability on the current trial but is largely insensitive to reliability on the previous trial. They termed this pattern “seemingly anti-Bayesian” because direct integration predicts that the uncertainty of both trials should matter, and related asymmetries have been reported in other paradigms (Ceylan et al., 2021; You et al., 2023). Experiment 1 showed an analogous pattern with motion coherence as the manipulation of stimulus quality: the previous-duration slope followed the current coherence, the previous coherence had no reliable effect, and the ordering predicted by static reliability weighting was not favored in either experiment. The three-state model accounts for the asymmetry as a consequence of sequential updating. What the preceding trial leaves behind, a shifted mean and the trace of its duration estimate, is already fixed when the current stimulus arrives. How much of the trace enters the prior is decided on the current trial, when the current coherence is known. The decomposition of the fitted slopes shows this directly (Figure 6C,F): the contribution of the retained trace depended on the current coherence and was the same after low- and high-coherence trials. A pattern that appears “anti-Bayesian” under direct integration can thus be accommodated by a multi-state updating architecture, without treating serial dependence as a processing error.

Whether stronger retention under low coherence benefits performance is a separate question. Retaining the previous trace reduces variability when the world is stable and successive durations are related (Cicchini et al., 2018; Kiyonaga et al., 2017; Manassi & Whitney, 2024), and it adds error when they are independent, as in the present sequences (Ozkirli et al., 2026). The account also complements models that explain the sign of history biases. Hahn and Wei (2024) reconciled seemingly contradictory attractive and repulsive biases within a Bayesian framework that accounts for encoding variability, and the observer of Fritsche et al. (2020) combines efficient encoding with Bayesian decoding to produce both biases concurrently. Whereas these models address the sign of the bias across estimation tasks, the three-state architecture separates the temporal routes through which prior information reaches the estimate.

### 6.3 Coherence transitions, perceptual continuity, and volatility

In Experiment 2, serial dependence was elevated when low coherence repeated, and the model reproduced the transition pattern only when the trace of the previous event was dropped after a change of coherence. Because abrupt onsets and a color change marked each encoding interval as a distinct perceptual event, this pattern suggests that carryover depends on the continuity between successive events. It resonates with event segmentation theory, in which the stream of experience is parsed at points of change and memory integration occurs preferentially within events (Baldassano et al., 2017; Zacks & Swallow, 2007). The evidence has limits. The planned Same/Switch interaction was not reliable, and the elevation was concentrated on LL trials and was identified in a follow-up contrast. The reset of the trace was added to the model space after this pattern had been observed, and the model with the reset predicts an elevation on HH trials as well, which was not observed. The reset is therefore best read as the simplest rule that reproduces the transition pattern. Retention that requires both weak current evidence and continuity with the preceding event would remove the remaining misfit, and it can be tested with a design that crosses the sharpness of event boundaries with coherence within participants.

Continuity between events should be distinguished from the volatility of the environment. In both experiments, coherence levels and durations were drawn independently on every trial with the same transition probabilities. A switch of coherence therefore carried no information about a change in the statistics of the durations, and the experiments do not test how observers respond to volatility. The model nevertheless makes definite predictions for such a manipulation, which we tested on the data of Glasauer and Shi (2022). With parameters fitted to the randomized order and then held fixed, the model predicted the near absence of central tendency and the reversal of the dependence on the previous duration in the random-walk order. In this framework, a change of sequence volatility alters behavior because the same observer encounters a differently ordered input: under a random walk the learned mean stays close to the current duration, so the pull toward it is small. Separate fits to the two orders showed no reliable difference in process noise or retention (Appendix C). Whether observers additionally adapt their updating to the volatility they experience over longer exposure remains open.

The temporal window analyses agree with a trace that is short-lived. In both experiments, serial dependence was confined to the few immediately preceding trials and decayed across lags, with no reliable future-trial effects, which indicates history dependence rather than slow drift (Section 4.3). Because the lag regressors were estimated in separate models, this conclusion should be treated with caution.

### 6.4 Response bias and limitations

Both experiments favored a bias that is added to the response (B2) over a bias that enters the observer’s prediction and over no bias state, by at least 15 AIC units. The bias decayed slowly (*λ* ≈ 0.72–0.78), so a response tendency persisted over several trials, in line with the dependence of the current error on the errors of the preceding responses. This response tendency is separable from the dependence on the previous stimulus: it produced correlated drifts of successive responses but did not contribute to the slope on the previous duration, and the coherence modulation of that slope remained when response history was controlled. Sequential effects in temporal reproduction thus have at least two sources, one tied to the history of the stimuli and one to the history of the responses. In the present model, the trace of the preceding trial enters the estimate that is reproduced, after the current stimulus has been processed, and the slow bias enters after it, in the response. This structure agrees with accounts that separate attraction to the previous response from the effect of the previous stimulus (Pascucci et al., 2019; Sadil et al., 2024), that place attractive biases at post-perceptual stages (Fritsche et al., 2017, 2020; Sheehan & Serences, 2022), and that tie the dependence to the internal state of the observer (Ozkirli & Pascucci, 2023). It is a property of the model, however, and behavioral fits do not identify a perceptual, decisional, or motor locus (Cicchini et al., 2024).

Several limitations remain. First, the reproduction display resembled the stimulus. The response-history analysis shows that the coherence effect is not carried by the previous responses, but reproducing with a moving-dot display may still raise the overall level of serial dependence. Second, the two experiments were run on separate samples and differed in several respects, so differences between them cannot be attributed to the sharpness of the event boundaries alone. Third, the model overestimates serial dependence on current-high trials (Section 5.4), assumes constant response noise, and does not identify where coherence acts on the observer’s noise, and individual retention values were recovered with limited precision (Appendix B.5–B.6). Fourth, the binary coherence manipulation does not capture the continuous variation in stimulus quality encountered in natural environments. Finally, the shorter reproductions under low coherence in Experiment 1 were modeled as an additive offset, and their likely origin is the ramped onset and offset of coherence. If observers register coherent motion when it passes a fixed criterion, the onset is registered later and the offset earlier when coherence is low, the more so because the low-coherence ramp was twice as long, and the interval is perceived as shorter by a fixed amount. This agrees with a shift that was the same at all durations and nearly absent in Experiment 2, in which coherence changed abruptly and a color change marked the interval. Because the two experiments differed in both respects, the account remains to be tested directly, by varying the ramp at a fixed coherence level.

### 6.5 Conclusion

Serial dependence in duration reproduction changed with motion coherence, whereas central tendency did not. With gradual onsets and offsets, serial dependence was stronger under low than under high current coherence. With sharply marked intervals, it was elevated when low coherence repeated. Within the tested Kalman-filter model space, a single precision-weighted observer predicted the consequences of a change in sequence volatility in independent data. More generally, the problem studied here arises wherever a changing world is estimated from noisy evidence: a system must decide how much of its recent past to carry into the present. Learners adjust the weight of new outcomes to the volatility of their environment (Behrens et al., 2007), artificial networks trained on tasks must protect earlier knowledge (Kirkpatrick et al., 2017), and raters who mark essays in sequence are drawn toward the score they have just given (Attali, 2011). For such systems, the present findings suggest that the use of the past has two separable parts. One is how strongly new evidence is weighted. The other is what the expectation contains: a long-run summary alone, or also a trace of the most recent event. In our data, stimulus quality and the continuity between events acted on the second part. Weak evidence can therefore lead a system to consult a prior while it trusts the evidence as before, and boundaries between events can mark where recent traces are dropped while long-run knowledge is kept. Whether this holds beyond one task and one model remains to be tested. A perceptual-timing task can thus illuminate how adaptive systems preserve useful continuity without letting outdated context dominate current inference.

## Data and Code Availability

Data and analysis code are available at https://github.com/msenselab/precision-weighting-serial-dependence.

## Appendix A: Linear Mixed Models and Behavioral Analyses

We report the primary trial-level LMMs and four coherence transitions (A.1), response-error history (A.2), mean reproduction and variability (A.3), and predicted slope orderings (A.4). Complete numerical summaries and the values plotted in the figures are available in the public repository (see Data and Code Availability).

### A.1 Duration History and Coherence Transitions

#### Primary inferential models

The primary LMMs modeled current reproduction bias as a function of centered current duration, centered previous duration, and the experiment-specific moderator. Models were fitted with restricted maximum likelihood and included participant random intercepts plus random slopes for current and previous duration:

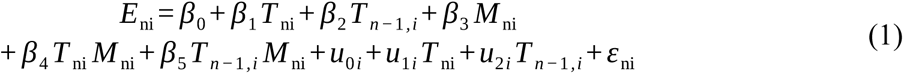

where E_ni_ is the reproduction bias on trial *n* for participant *i* (reproduced minus presented duration); *T*_ni_ and *T_n−_*_1*,i*_ are the centered current and previous durations; *M*_ni_ is the experiment-specific moderator (current low coherence in Experiment 1; Same/Switch transition in Experiment 2); *u*_0*i*_ *, u*_1*i*_ *, u*_2*i*_ are participant-level random intercepts and slopes; and *ε*_ni_ is the residual. Fixed-effect estimates for the moderator interactions are shown in Figure A1A.

These are the models underlying the main text claims. In both experiments, the current duration × moderator interaction was small and non-significant, providing no evidence that central tendency was modulated by current coherence (Experiment 1) or by Same/Switch transition status (Experiment 2). The previous duration × moderator interaction was reliable in Experiment 1, showing that the controlled serial dependence channel was selectively strengthened by low current coherence, and was not reliable in Experiment 2, where the descriptive Same > Switch pattern was weaker at the trial level.

### Four coherence transitions

To compare the same four transitions in both experiments, a common model crossed previous and current coherence. Both durations were centered at 1.2 s, and the fixed effects were current duration × previous coherence × current coherence + previous duration × previous coherence × current coherence, with participant random intercepts and random slopes for current and previous duration (4,693 and 4,654 trials, 22 participants per experiment). Figure A1B–D shows the previous-duration slopes and their planned and post-hoc contrasts. Transitions are labeled with the previous coherence first.

**Figure A1:**
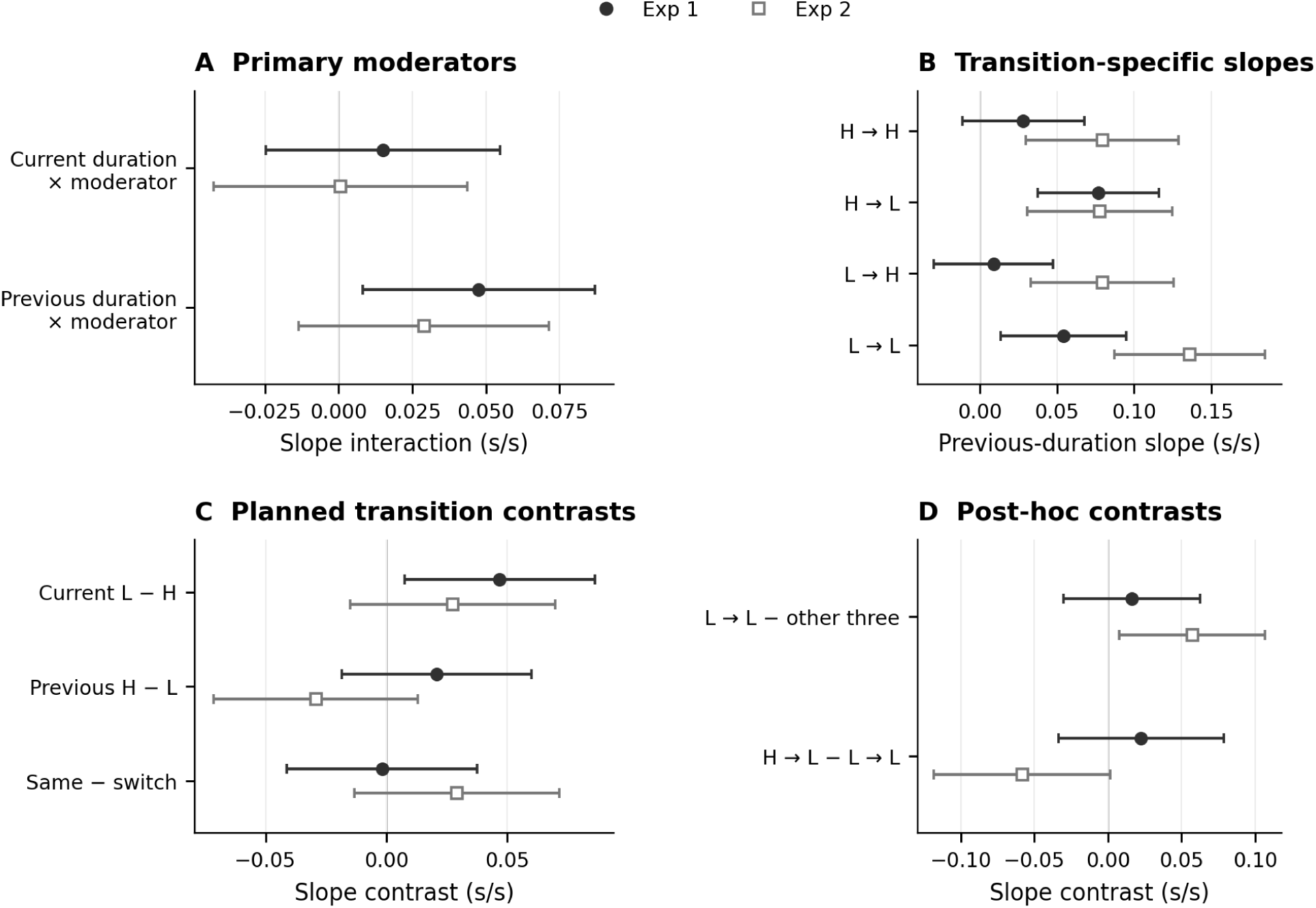
Trial-Level Duration-History Effects and Coherence-Transition Contrasts. *Note.* Points and bars are the mixed-model estimates and pointwise 95% confidence intervals. Filled circles denote Experiment 1 and open squares Experiment 2. (A) Primary interaction estimates: the moderator is current Low − High coherence in Experiment 1 but Same − Switch transition in Experiment 2; these are different contrasts, not an experiment interaction. (B) Controlled previous-duration slopes; transition labels give previous then current coherence. (C) Three planned contrasts of those slopes. Their raw/Holm-adjusted *p* pairs, in row order, are .020/.059, .298/.597, and .931/.931 in Experiment 1 and .203/.529, .176/.529, and .177/.529 in Experiment 2. (D) Four post-hoc contrasts, shown with nominal intervals and *p* values; LL minus the other three in Experiment 2 has nominal *p* = .024 and four-follow-up Bonferroni *p* = .096. All intervals are unadjusted; neither interval overlap nor exclusion of zero replaces the stated correction families.

The current-coherence contrast in Experiment 1 was similar across random-effects specifications (intercept only; with either or both duration slopes) and with participant-clustered standard errors (estimates 0.046–0.047, *p* = .006–.028). The Same − Switch estimate in Experiment 2 was positive in all specifications (0.019–0.029, *p* = .166–.197).

### A.2 Response-Error History

The reproduction display resembled the stimulus, and each response may leave a trace of its own. To separate stimulus history from response history, we defined the response error of a preceding trial as the reproduced minus the presented duration and entered the errors of the five preceding trials together with current and previous physical duration:

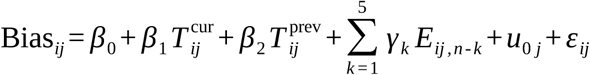

Lagged errors were constructed within participant and block on the complete trial sequence, before response exclusions (2,802 and 2,659 trials with complete histories). Response-error carryover was strongest at lag 1 and decayed across lags in both experiments (Figure A2A), consistent with a gradually decaying response-bias state. Variance inflation factors were modest (largest VIF = 1.57 and 1.46), so the lagged-error model did not suffer from problematic multicollinearity.

**Figure A2:**
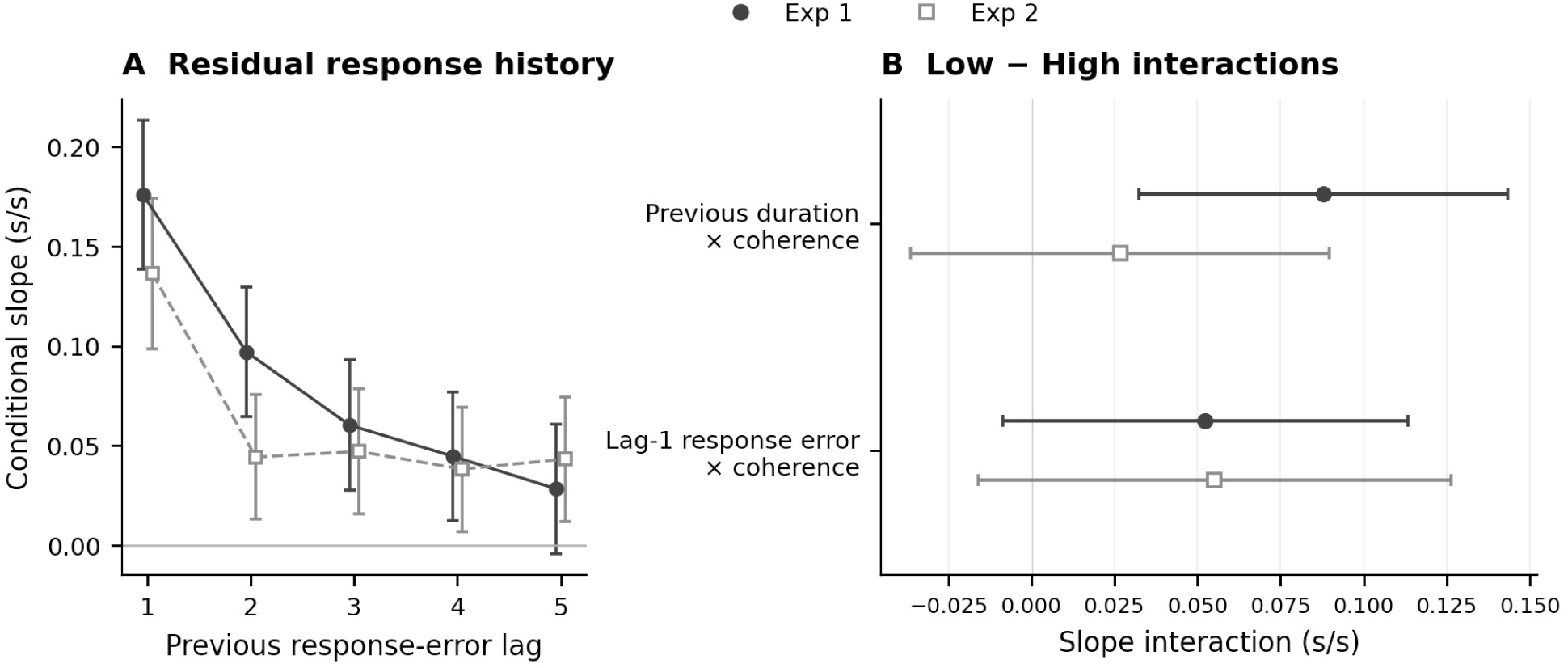
Response-Error Carryover and Coherence-Dependent History Effects. *Note.* Filled circles/solid lines denote Experiment 1; open squares/dashed lines Experiment 2. Bars are pointwise 95% confidence intervals. (A) Response-error lags 1–5 entered simultaneously, controlling current and previous duration (2,802 and 2,659 complete-history trials). (B) Low − High slope interactions from the separate coherence-interaction model, with response-error lags 2–5 as covariates. Both models include participant random intercepts. Lag tests are raw; the two coherence interactions form a separate Holm family within each experiment. No significance coding is added.

In Experiment 1, the coherence modulation of the previous-duration slope remained when response history was included (Holm-adjusted over the two interactions, *p* = .004). The same interaction estimated participant by participant was in the same direction and less conclusive (mean *b* = 0.074, *t*(21) = 1.91, *p* = .070). These analyses separate the physical duration of the preceding trial from the residual history of responding. They do not assign either effect to a perceptual, decisional, or motor stage.

### A.3 Mean Reproduction, Response Variability, and Design Sensitivity

#### Mean shift

The current Low − High contrast in mean reproduction was estimated in each experiment from the four-transition model at current and previous durations of 1.2 s, averaging equally over previous coherence. The Low − High difference was −0.084 s in Experiment 1 (*SE* = 0.006, 95% CI [−0.095, −0.073], *p* < .001) and −0.012 s in Experiment 2 (*SE* = 0.006, 95% CI [−0.024, 0.000], *p* = .052). In a combined model with experiment as a factor, the two differed by −0.073 s (95% CI [−0.089, −0.056], *p* < .001). Participant-level estimates gave the same pattern (Experiment 1: −0.083 s, *t*(21) = −5.10, *p* < .001; Experiment 2: −0.011 s, *t*(21) = −1.29, *p* = .212).

To see whether the shift in Experiment 1 was additive or proportional to the duration (Figure A3A), the Low − High difference of the mean reproductions was computed for each participant at each duration. It was −0.083, −0.087, −0.088, −0.084, and −0.072 s at 0.8, 1.0, 1.2, 1.4, and 1.6 s (*SEM* between 0.019 and 0.024 s). The participant-level slope of the shift on duration did not differ from zero (0.011, *SE* = 0.028, *t*(21) = 0.41, *p* = .685). It differed from the slope expected if low coherence shortened all durations by the same proportion (−0.069; *t*(21) = 2.88, *p* = .009).

#### Response variability

The SD of the reproductions was computed within each participant, physical duration, and coherence level, and averaged over the five durations (Figure A3B). Mean Low/High SDs were 0.193/0.173 s in Experiment 1 and 0.206/0.201 s in Experiment 2; the Low − High contrasts were 0.021 s (95% CI [0.011, 0.031], *p* < .001) and 0.005 s (95% CI [−0.002, 0.013], *p* = .152). The median absolute deviation gave the same result (Experiment 1: 0.012 s, *p* = .014; Experiment 2: −0.001 s, *p* = .895).

#### Design sensitivity

The a priori power analysis targeted the basic within-participant serial dependence effect (*d*z = 0.70), not its moderation by coherence or transition. With *N* = 22, a two-sided paired test at alpha = .05 has 80% power for *d*z ≥ 0.63. The meta-analysis of Manassi et al. (2023) provides effect sizes for serial dependence in visual tasks; it does not provide an effect size for the moderation of serial dependence in duration reproduction by stimulus quality, so we report the sensitivity of the design instead of a meta-analytic power estimate. The observed moderation in Experiment 1 was *d*z = 0.75.

### A.4 Comparison of Predicted Orderings

The three accounts in the Introduction predict different orderings of the four slopes (Section 3.3). We compared them using one controlled previous-duration slope per participant and transition. Within an ordering, two transitions counted as approximately equal when they differed by no more than δ, and as ordered when they differed by more than *g*. The reported values (Figure A3C) use δ = *g* = .02; a slope difference of .02 corresponds to 16 ms across the 0.8–1.6 s stimulus range. Bayes factors against unrestricted means were computed with the encompassing-prior approach (Hoijtink et al., 2019): the posterior probability of the constrained region divided by its prior probability, under a multivariate normal model of the participants’ contrast vectors with a conjugate normal–inverse-Wishart prior (prior scale τ = .10; Murphy, 2007). The bounds and the LL-elevation ordering were specified for this follow-up analysis.

Grouping by current coherence was favored over static reliability weighting in Experiment 1 (ratio of Bayes factors = 6.4), and the elevation on LL was favored over grouping by current coherence in Experiment 2 (60.8). Across three equality bounds, two gap definitions, and three prior scales (18 settings), the corresponding ranges were 1.9–15.1 and 7.9–139.9, and the first-ranked ordering was the same in all settings and in all 22 leave-one-participant-out analyses of each experiment. The Bayes factors compare patterns of slopes; they are not probabilities of cognitive mechanisms.

**Figure A3:**
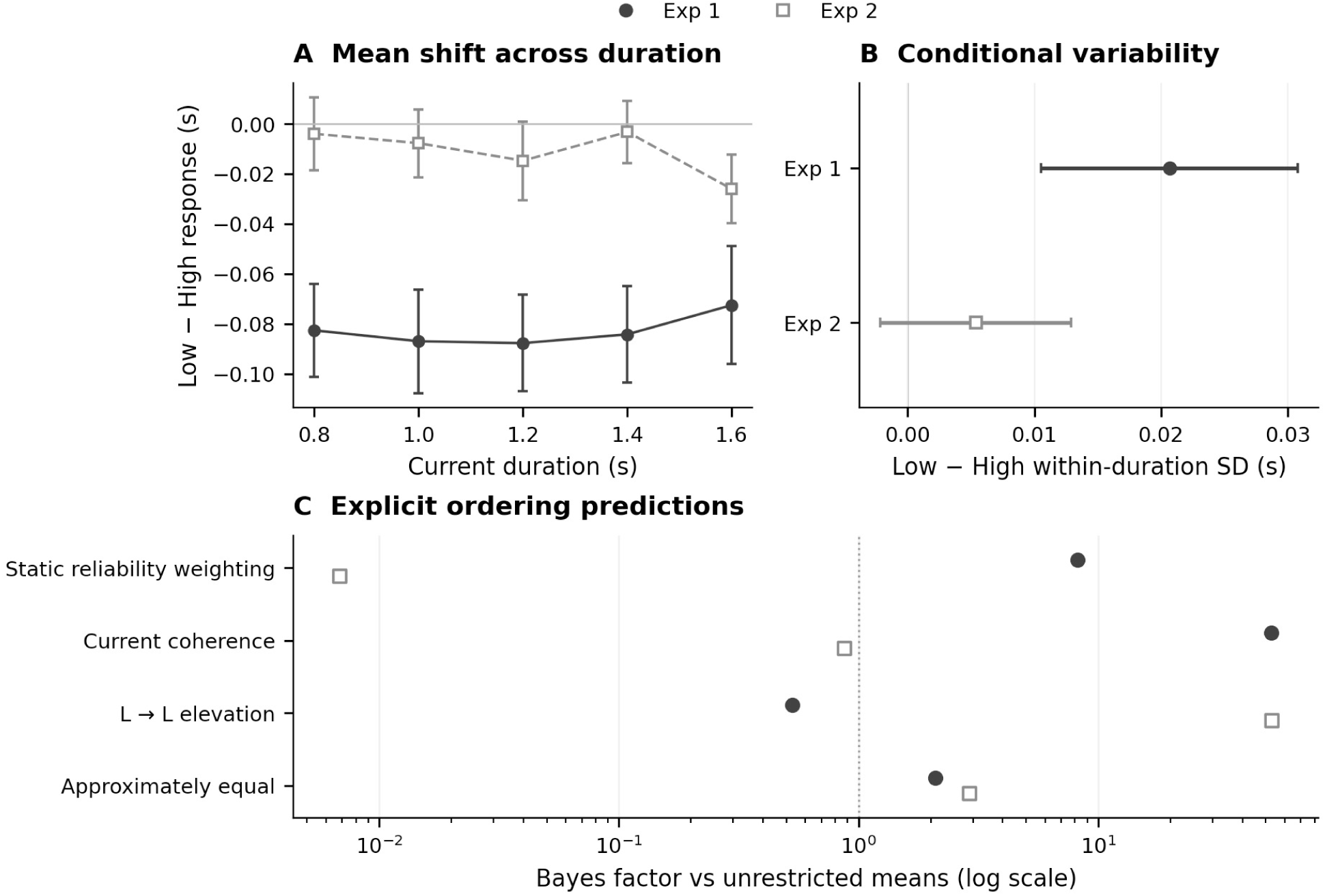
Mean Reproduction, Conditional Variability, and Evidence for Slope Orderings. *Note.* Filled circles/solid lines denote Experiment 1; open squares/dashed lines Experiment 2. (A) Participant mean Low − High reproduction at each physical duration ± SEM; these descriptive cell means are distinct from the adjusted LMM offsets in the text. The additive-versus-proportional tests use participant slopes, not independent duration cells. (B) Within-duration reproduction-SD contrasts with paired 95% confidence intervals; this is not the residual SD of the model’s predictive check. (C) Bayes factors against unrestricted slope means on a log scale, at τ = .10 and δ = *g* = .02. The tested patterns are static reliability, HL > {LL, HH} > LH; current coherence, {HL, LL} > {HH, LH}; LL elevation, LL > {HH, LH, HL}; and approximate equality of all four transitions. Braces denote approximate equality. The reference at 1 is equal evidence against the unrestricted comparison, not a significance threshold. These are separate estimands and analyses, not a combined test.

## Appendix B: Computational Modeling Technical Details

### B.1 The Three-State Kalman Filter

#### States

The observer’s state on trial *i* is **s***_i_*=_[_*x̂_i_, m̂_i_, b̂_i_*_]_^⊤^, where *x̂* is the fast state (the current duration estimate), *m̂* the slow state (the learned mean), and *b̂* the bias state. 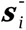 denotes the prediction and **s***_i_* the posterior. *s_i_* is the presented duration.

#### State evolution

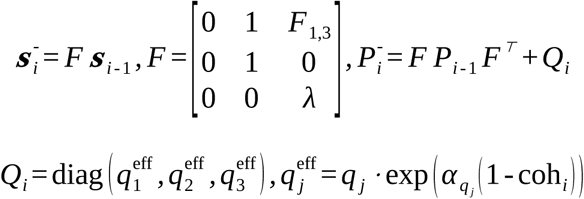

The modulation 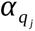 is free only for the noise terms selected by the C level and is zero otherwise. The first row of *F* gives the prediction of the fast state, 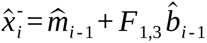: the prediction starts from the learned mean, and the fast state of the preceding trial is not carried into the filter. The learned mean and the bias evolve as 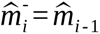 and 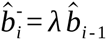.

#### Observation and update

*H* = [1,0,0]. The measurement is the presented duration, *z_i_* = *s_i_*; the measurement noise *R_i_* expresses the reliability that the observer assigns to it and sets the gain.

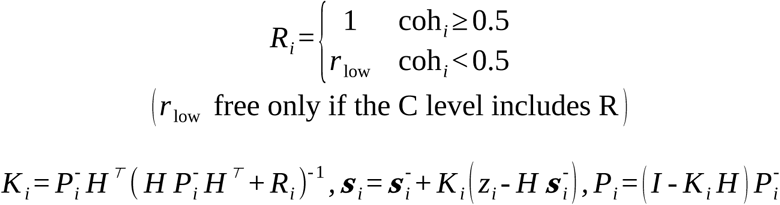

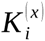 and 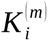 denote the components of *K_i_* for the fast and the slow state.

#### Readout

With the fast deviation *f̂_i_* = *x̂_i_* - *m̂_i_*, the readout is

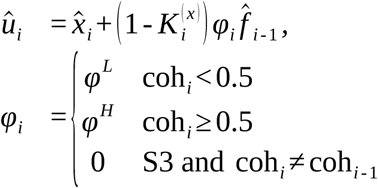

This is the posterior mean under the readout prior 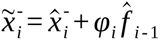 with the gain of the filter: 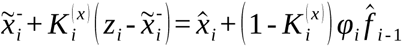. Only (**s***_i_, P_i_*) are carried to the next trial, so the readout does not feed back into the filter. With *φ* = 0 the readout is *x̂_i_*. For models without a bias in the prediction 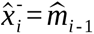 and the readout prior equals (1 - *φ_i_*) *m̂_i-1_*+ *φ_i_x̂_i-1_*, the form used in Equation 2 of the main text. The trace of the main text is the previous estimate *x̂_i_* _- 1_, and the retention *φ_i_* is its weight in this prior.

#### Response

With the offset 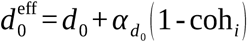 and response noise 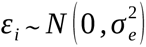

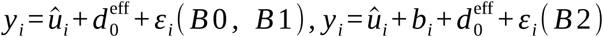

Under B2, *b_i_*= *λ b_i_*_-1_ + *η_i_* with 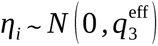 is a latent response bias that the observer does not track (*F*_1,3_ = 0); the likelihood integrates over it.

### B.2 Central Tendency and Serial Dependence in the Model

For models without a bias in the prediction (B0, B2) and without a switch effect (S0), the update is 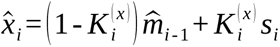 and 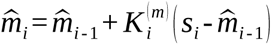, so that 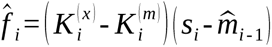. Inserting these into the readout gives the slope of the readout on the current duration,

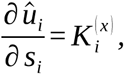

which does not depend on *φ*, and the slope on the previous duration,

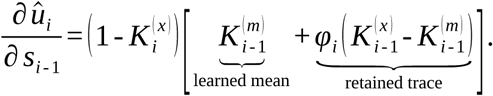

Central tendency is therefore set by the gain alone. Serial dependence has two routes: the previous duration moved the learned mean, and a share *φ_i_* of the previous fast deviation is retained. The current coherence can change serial dependence through the factor 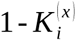, if coherence changes the gain, or through *φ^L^ ≠ φ^H^*. The gains have the closed form 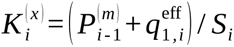 and 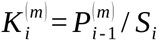 with 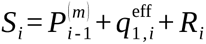, where *P*^(*m*)^ is the uncertainty of the learned mean; hence 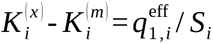. Under S1 the first term in the bracket is multiplied by 1 - *γ* on switch trials, and under B1 the term 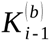 is added. These expressions agree with the implemented model to numerical precision. Figure 6C,F of the main text shows the two terms of the bracket, multiplied by 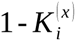, at the fitted parameters.

### B.3 Model Space, Fitting, and Comparison

Every model combines one level of each of three dimensions (Table B1).

**Table B1:**
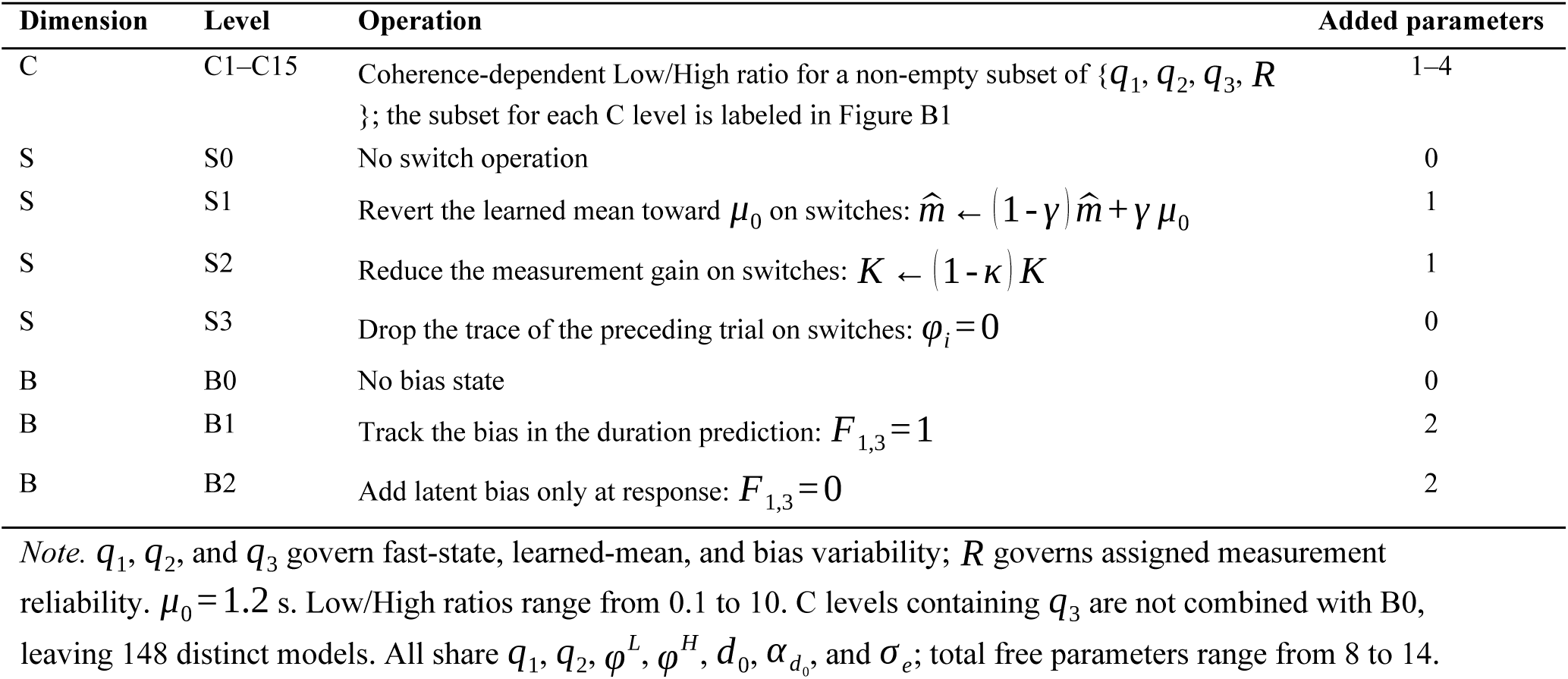
Operations Defining the Three Model Dimensions.

| Dimension | Level | Operation | Added parameters |
|---|---|---|---|
| C | C1–C15 | Coherence-dependent Low/High ratio for a non-empty subset of {*q*<sub>1</sub>, *q*<sub>2</sub>, *q*<sub>3</sub>, *R* }; the subset for each C level is labeled in Figure B1 | 1–4 |
| S | S0 | No switch operation | 0 |
| S | S1 | Revert the learned mean toward *μ*<sub>0</sub> on switches: *m̂* ← (1 - *γ*) *m̂* + *γ μ*<sub>0</sub> | 1 |
| S | S2 | Reduce the measurement gain on switches: *K* ← (1 - *κ*) *K* | 1 |
| S | S3 | Drop the trace of the preceding trial on switches: *φ*<sub>*i*</sub> = 0 | 0 |
| B | B0 | No bias state | 0 |
| B | B1 | Track the bias in the duration prediction: *F*<sub>1,3</sub> = 1 | 2 |
| B | B2 | Add latent bias only at response: *F*<sub>1,3</sub> = 0 | 2 |

*Note.* *q*<sub>1</sub>, *q*<sub>2</sub>, and *q*<sub>3</sub> govern fast-state, learned-mean, and bias variability; *R* governs assigned measurement reliability. *μ*<sub>0</sub> = 1.2 s. Low/High ratios range from 0.1 to 10. C levels containing *q*<sub>3</sub> are not combined with B0, leaving 148 distinct models. All share *q*<sub>1</sub>, *q*<sub>2</sub>, *φ*<sup>*L*</sup>, *φ*<sup>*H*</sup>, *d*<sub>0</sub>, *α*<sub>*d*<sub>0</sub></sub>, and *σ*<sub>*e*</sub>; total free parameters range from 8 to 14.

#### Fitting

Models were fitted to the trial-wise presented and reproduced durations of each participant (22 per experiment) by maximum likelihood on the linear duration scale. Because the model is linear and Gaussian, the likelihood of the responses is exact; under B2 the latent response bias is integrated out with a Kalman filter over the response residuals. Trials excluded in preprocessing were presented to the observer but did not contribute to the likelihood. The learned mean was initialized at 1.2 s with a large variance, so that the first duration determines it.

Each fit used 12 starting points within the parameter bounds and one additional start with *φ^L^* = *φ^H^* = 0. Bounds were set on the natural scale of each parameter: *φ^L^*, *φ^H^* in [0, 1]; *λ* in [0, 0.99]; the response-noise SD *σ_e_* in [0.01, 0.5] s; the offset *d*_0_ and its Low − High difference in [−0.5, 0.5] s; *q*_1_ and *q*_2_ in [0, 25] in units of the high-coherence measurement noise; the SD of the response-bias drift in [0, 0.5] s; and every Low/High ratio in [0.1, 10]. Model comparison used *AIC* = 2 *k* - 2 *l* (Burnham & Anderson, 2002), averaged over participants.

#### Screening by simulation

Each fitted model was simulated 20 times on each participant’s trial sequence. For each statistic (central tendency and serial dependence by current coherence and by transition, and the residual SD), the model mean was compared with the data mean in units of the standard error of the data; a statistic counted as reproduced when the absolute difference was below 1.5.

#### Results

Table 1 of the main text summarizes the bias and switch dimensions. Figure B1 compares the 15 C levels in combination with the selected switch and bias levels. Differences in AIC between C levels were small, and the gain on the current duration was nearly the same under low and high coherence at every level.

**Figure B1:**
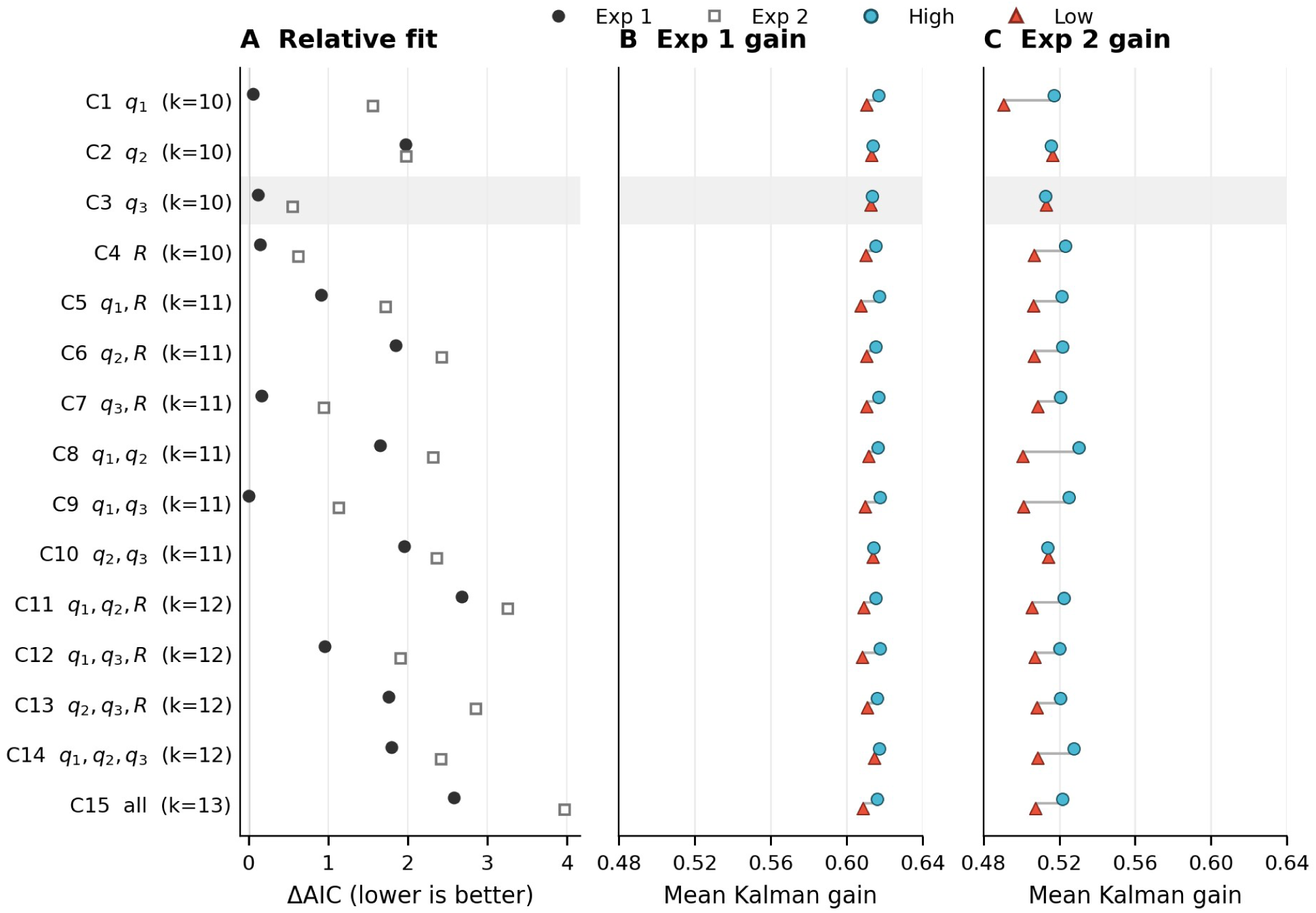
Coherence-Dependent Noise: Relative Fit and Low/High Measurement Gain. *Note.* (A) Descriptive forest-style point estimates of mean participant ΔAIC, relative to the best of the 148 models within each experiment; filled circles denote Experiment 1 and open squares Experiment 2. No confidence intervals are implied or added. Each C level is combined with S0/B2 in Experiment 1 and S3/B2 in Experiment 2. (B–C) Mean *_K_*^(^*^x^*^)^ on current High (cyan circles) and Low (red triangles) trials, averaging within participant then across the 22 participants. Small vertical offsets distinguish overlapping markers; gray segments indicate the Low/High span, not uncertainty. The two gain panels share a scale. The shaded C3 row identifies the coherence dimension used for the retained models; the AIC minimum need not be C3. *k* is the number of free parameters.

#### Parameter estimates

Figure 6 of the main text shows the gains and the retention of the selected models. Table B2 gives their remaining parameters.

**Table B2:**
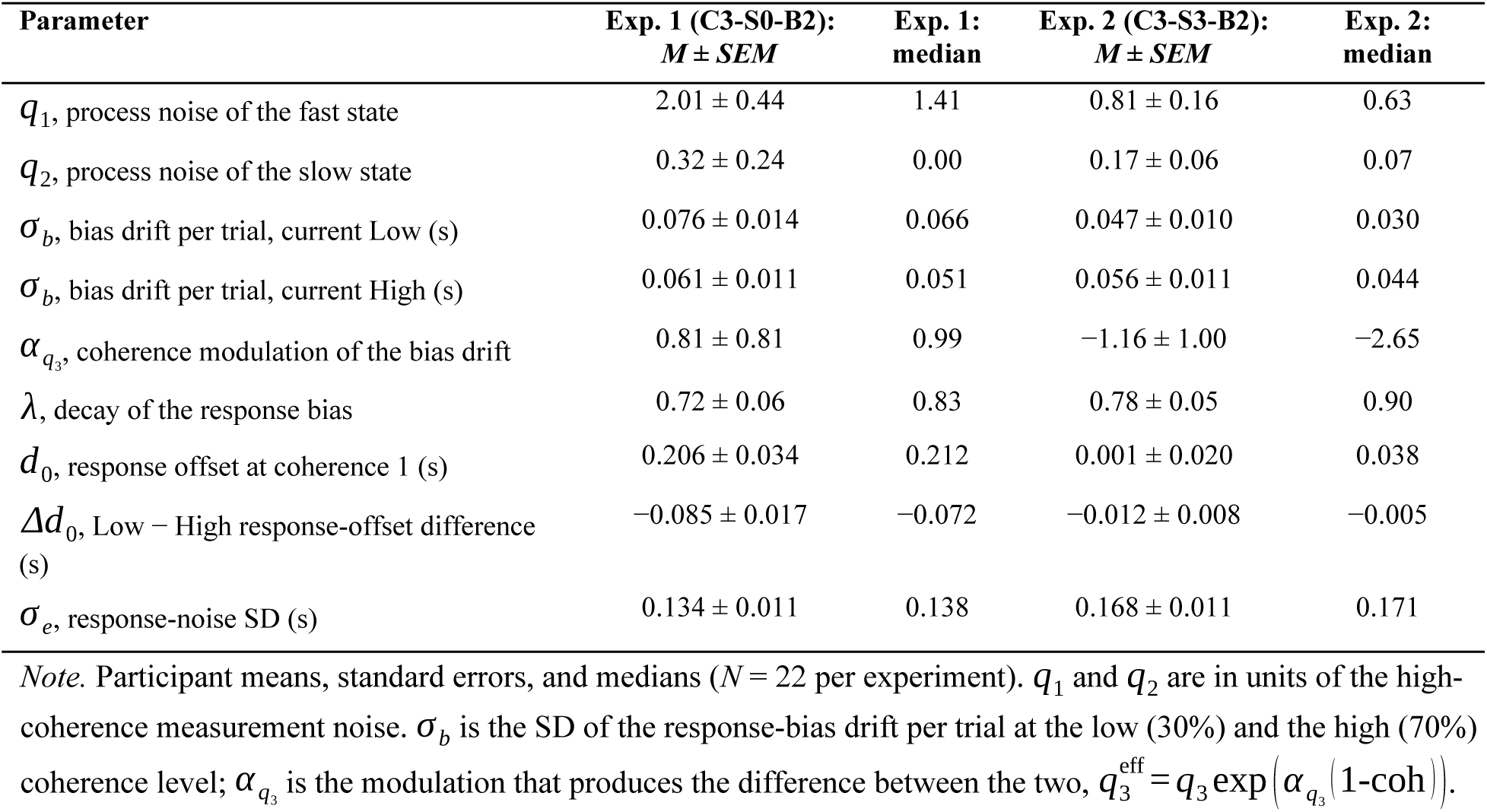
Remaining Parameter Estimates of the Selected Models.

| **Parameter** | **Exp. 1 (C3-S0-B2):<br>*M* ± *SEM*** | **Exp. 1:<br>median** | **Exp. 2 (C3-S3-B2):<br>*M* ± *SEM*** | **Exp. 2:<br>median** |
|---|---|---|---|---|
| *q*<sub>1</sub>, process noise of the fast state | 2.01 ± 0.44 | 1.41 | 0.81 ± 0.16 | 0.63 |
| *q*<sub>2</sub>, process noise of the slow state | 0.32 ± 0.24 | 0.00 | 0.17 ± 0.06 | 0.07 |
| *σ*<sub>*b*</sub>, bias drift per trial, current Low (s) | 0.076 ± 0.014 | 0.066 | 0.047 ± 0.010 | 0.030 |
| *σ*<sub>*b*</sub>, bias drift per trial, current High (s) | 0.061 ± 0.011 | 0.051 | 0.056 ± 0.011 | 0.044 |
| *α*<sub>*q*<sub>3</sub></sub>, coherence modulation of the bias drift | 0.81 ± 0.81 | 0.99 | −1.16 ± 1.00 | −2.65 |
| *λ*, decay of the response bias | 0.72 ± 0.06 | 0.83 | 0.78 ± 0.05 | 0.90 |
| *d*<sub>0</sub>, response offset at coherence 1 (s) | 0.206 ± 0.034 | 0.212 | 0.001 ± 0.020 | 0.038 |
| *Δd*<sub>0</sub>, Low − High response-offset difference (s) | −0.085 ± 0.017 | −0.072 | −0.012 ± 0.008 | −0.005 |
| *σ*<sub>*e*</sub>, response-noise SD (s) | 0.134 ± 0.011 | 0.138 | 0.168 ± 0.011 | 0.171 |

*Note.* Participant means, standard errors, and medians (*N* = 22 per experiment). *q*<sub>1</sub> and *q*<sub>2</sub> are in units of the high-coherence measurement noise. *σ*<sub>*b*</sub> is the SD of the response-bias drift per trial at the low (30%) and the high (70%) coherence level; *α*<sub>*q*<sub>3</sub></sub> is the modulation that produces the difference between the two, *q*<sub>3</sub><sup>eff</sup> = *q*<sub>3</sub> exp(*α*<sub>*q*<sub>3</sub></sub>(1-coh)).

### B.4 Predictive Checks by Duration

For the predictive checks, 100 cohorts were simulated from the fitted parameters; in each cohort every participant was simulated once on their own trial sequence. Figure B2 overlays Low and High coherence within each duration-resolved check, with experiments in adjacent columns. Reproduction bias by current duration and by previous duration was reproduced in both experiments. The residual SD of the model was flat across durations. In Experiment 1 the observed residual SD increased with duration (from about 0.18 to 0.20 s under low and from 0.16 to 0.19 s under high coherence), so that the model was too high at short and too low at long durations; Experiment 2 showed no clear trend.

**Figure B2:**
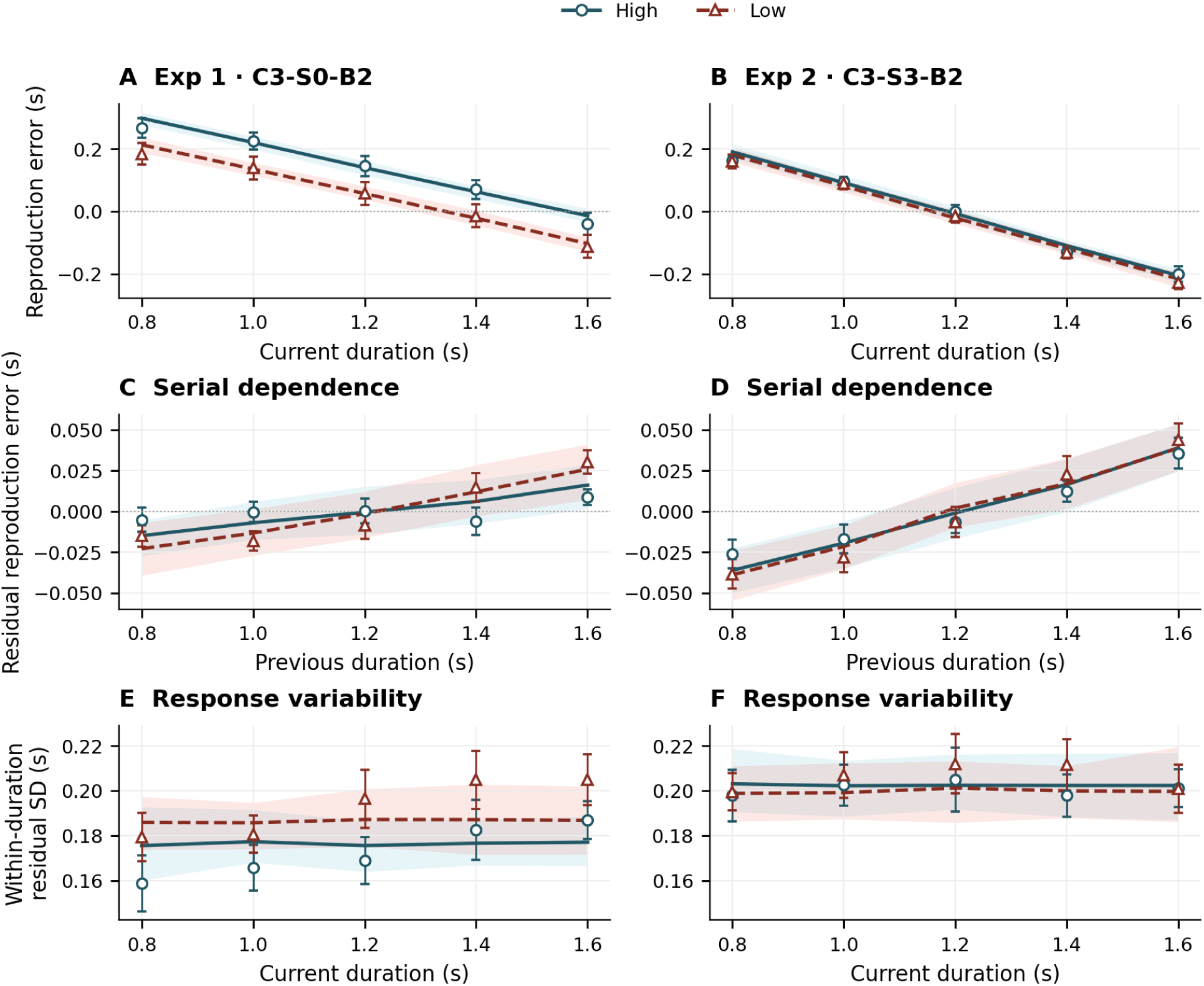
Duration-Resolved Predictive Checks With Low and High Coherence Overlaid. *Note.* Columns show Experiments 1 (C3-S0-B2) and 2 (C3-S3-B2); rows show bias by current duration, residual bias by previous duration, and within-duration residual SD. High coherence uses cyan circles and solid model lines; Low coherence uses red triangles and dashed lines. Open points are participant means ± SEM; model lines are means and shaded bands the central 95% range of cohort means across 100 simulated cohorts. Bands are predictive ranges, not confidence intervals for the observed means. Within each row, both experiments share a y-scale. Residual bias removes the participant/coherence/current-duration mean and is not the controlled serial-dependence slope of the LMM.

### B.5 Parameter and Coherence-Effect Recovery

To assess which fitted quantities can be estimated under the experimental design, we generated responses from the selected models, C3-S0-B2 in Experiment 1 and C3-S3-B2 in Experiment 2. Each of the 22 fitted participants per experiment generated 20 independent response datasets on their own 240-trial duration/coherence sequence, retaining the original response-validity mask. The observer processed the full sequence, including trials without a scored response. Newly drawn latent response bias and response noise varied across repetitions, and all ten free parameters were jointly re-estimated. The primary design therefore contained 880 simulated datasets. A further 400 exploratory parameter vectors sampled the fitted ranges, with continuity varying independently over 0–1; these results are available in the public repository.

The ten recovered quantities were the innovation SDs of the fast, slow, and bias states (the square roots of the process-noise variances); Low and High readout continuity (the retention parameters *φ^L^*and *φ^H^* of the main text); bias persistence; the baseline response offset and its Low − High difference; response-noise SD; and the Low/High bias innovation variance ratio. In the implementation, innovation SDs are optimized in seconds; the filter uses normalized process variances 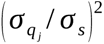, with *σ_s_* = 0.1 fixed in these selected models. The baseline offset and bias innovation SD are defined at coherence 1, rather than at the High condition. Condition offsets are *d _H_* = *d*_0_ + 0.75 *Δd*_0_ and *d _L_* = *d*_0_ + 1.75 *Δd*_0_. Both selected C3 models have *R_H_* = *R_L_* = 1.

Initial fits used 13 starting points: the default point, 11 random points, and a zero-continuity point. Fits that did not converge or fell more than 0.01 log-likelihood below the generating parameters received 49 additional starts, without using the generating parameters as an optimization start. Original and refined outputs were retained separately, and refinements used identical simulated responses. All 880 primary selected fits converged and met the generating-likelihood check after refinement. Figure B3 distinguishes recovery from one dataset per person from recovery after averaging 20 estimates per person. Full bias, RMSE, and boundary summaries are available in the public repository.

Fast innovation SD and response offsets recovered more consistently than individual readout continuity. For High continuity, single-cohort correlations were .157 and .258 in Experiments 1 and 2, compared with .663 and .690 after averaging each participant’s 20 estimates. High continuity reached its lower bound in 46.8% and 35.7% of recoveries. Thus, the parameter estimates do not support precise individual differences in High continuity. Figure B3 compares both recovery summaries for all ten fitted parameters; averaging repeated datasets must not be read as recovery from a single experiment.

**Figure B3:**
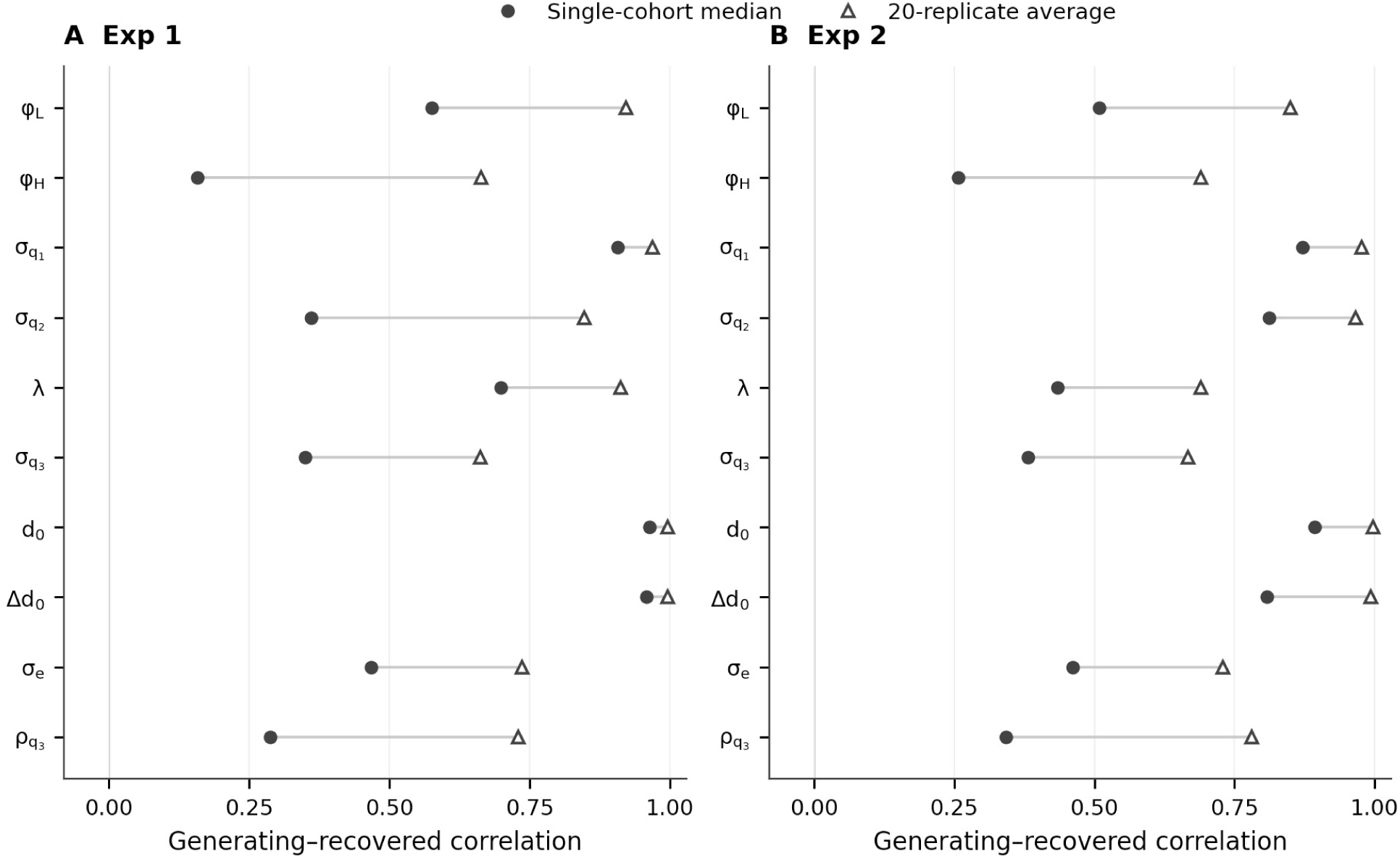
Single-Dataset Versus Repeated-Dataset Parameter Recovery. *Note.* Filled circles show the median generating–recovered correlation over 20 cohorts, each with one dataset for each of 22 participants. Open triangles show the correlation of the 22 generating values with participant estimates averaged over 20 datasets. Connecting segments distinguish these two summaries and are not confidence intervals. The two experiments share the correlation scale. 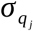 are innovation SDs; *λ* is bias persistence; *d*_0_ and *Δd*_0_ are baseline and Low − High offsets; *σ_e_* is response noise; and 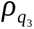 is the bias-variance ratio. The weak single-dataset recovery of *φ^H^* remains visible rather than being obscured by the higher repeated-dataset correlations.

We also recovered the Low − High continuity difference, *Δφ* = *φ^L^* - *φ^H^*, and the corresponding difference in effective trace weight, 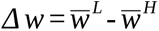. Trial weights were computed as 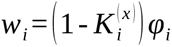 and averaged separately over Low and High trials on the actual sequence. Under S3, switch trials contribute zero trace weight. These readout weights are distinct from the complete slope on the preceding duration, which also includes the learned-mean route (B.2). The mean generating effects were positive in both experiments, but their direction recovered more consistently in Experiment 1 (Figure B4).

**Figure B4:**
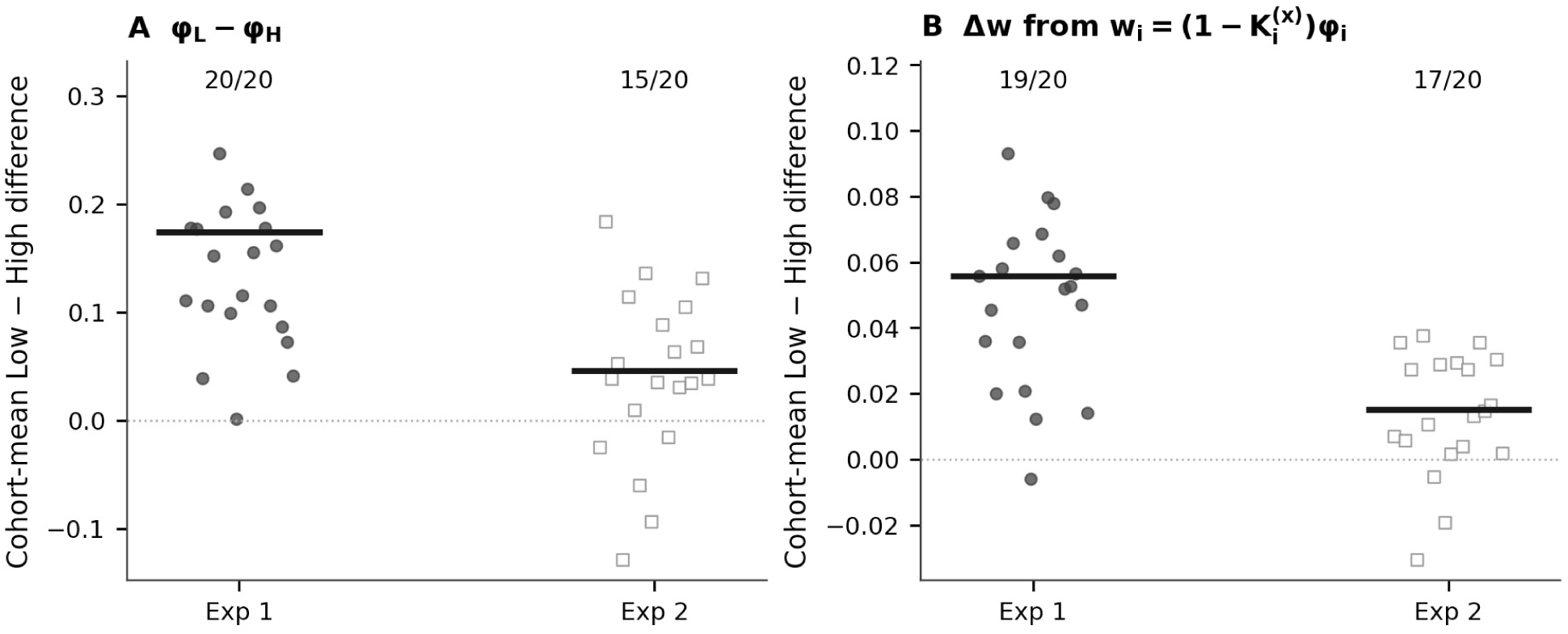
Recovery of Cohort-Mean Coherence Differences. *Note.* Each point is one recovered 22-participant cohort mean, with 20 independent response cohorts per experiment; filled circles denote Experiment 1 and open squares Experiment 2. Horizontal offsets only separate repetitions. Short thick horizontal lines show the fixed generating cohort means. Panels show (A) *Δφ* and (B) *Δw*, with 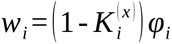 averaged within current coherence; fractions give the number of recovered cohort means with the generating sign (20/20 and 15/20 for continuity; 19/20 and 17/20 for trace weight). These are conditional recovery frequencies at the fitted generating values, not empirical-effect tests or estimates of design power.

### B.6 Model Recovery and High-Continuity Identifiability

#### S0 versus S3

To test discrimination of the switch rule, each generating structure produced 100 cohorts on the Experiment 2 sequences, with 22 simulated participants per cohort. Every simulated participant dataset was fitted with both C3-S0-B2 and C3-S3-B2, giving 8,800 candidate fits. Both candidates have ten free parameters. Within each cohort we selected the candidate with the lower mean participant AIC. S0 was selected in 99 of 100 S0-generated cohorts, and S3 in 91 of 100 S3-generated cohorts (Figure B5). At the individual-dataset level, correct selections were 1,429 of 2,200 for S0 and 1,312 of 2,200 for S3. The strong cohort-level discrimination therefore does not imply reliable classification of individual participants. In the observed data of Experiment 2, the mean AIC of C3-S0-B2 was lower than that of C3-S3-B2 by 0.56 units. A difference of at least this size in favor of S0 occurred in 83 of 100 S0-generated cohorts and in 2 of 100 S3-generated cohorts. By AIC alone, the observed cohort therefore resembles the S0-generated cohorts, although the S0 models do not reproduce the elevation on LL trials (Table 1 of the main text). These simulations compare this pair at fitted generating values on Experiment 2 schedules; they do not identify the C dimension, test the entire model family, or establish different mechanisms between experiments. All cross-fit AIC values are finite, but optimizer-success and generating-likelihood checks were not recorded for these fits, so the convergence statement in B.5 does not extend to them.

#### High-continuity profiles

For each of the 44 observed participants, we fixed High readout continuity at values from 0 to 1 and reoptimized the other nine parameters with multiple starts and a reverse warm-start pass. An unrestricted refinement checked the reference optimum; no restricted point exceeded that retained optimum. Figure B5B–C uses twice the log-likelihood loss of 3.84 as a descriptive reference, not a calibrated confidence boundary. The whole 0–1 grid lay below it for 4 of 22 participants in Experiment 1 and 16 of 22 in Experiment 2; median supported grid spans were 0.5 and 1.0. The S3 rule in Experiment 2 reduced the median number of usable High trials with an active continuity path to 49.5, compared with 107 in Experiment 1. Two Experiment 2 fits had zero fast innovation SD, making the fast deviation zero and continuity irrelevant to the predicted response. One of these flat, finite profiles had failed optimizer flags at the unrestricted solution and its 12 grid points, despite matching the retained likelihood. These profiles agree with the recovery results: individual High continuity is often weakly identified, especially in Experiment 2.

**Figure B5:**
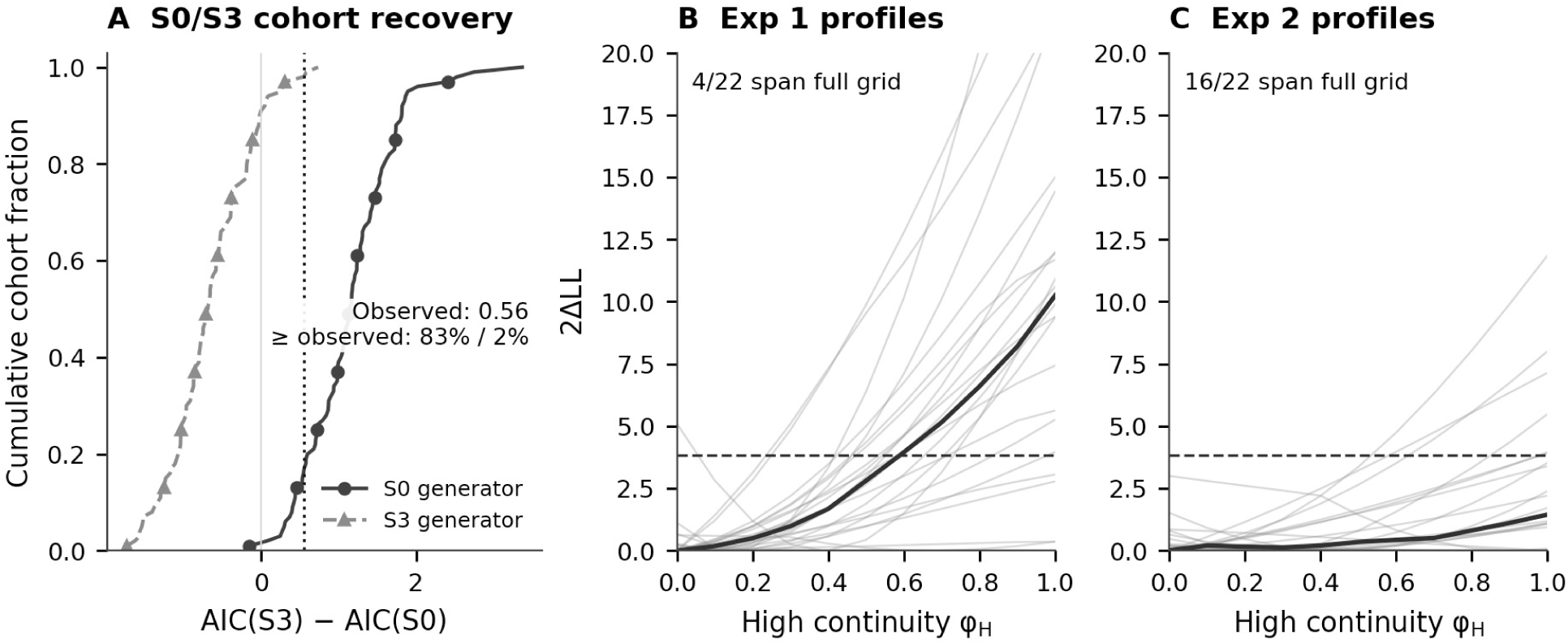
Cohort-Level Switch-Rule Discrimination and Individual High-Continuity Profiles. *Note.* (A) Empirical cumulative distributions of mean AIC(S3) − AIC(S0) for 100 cohorts per generating rule on Experiment 2 schedules. Solid circles denote S0-generated cohorts and dashed triangles S3-generated cohorts. Positive values favor S0; the dotted line marks the observed difference of 0.56. The 83%/2% annotation gives fractions at least that large under S0/S3 generation, not *p* values. (B–C) Thin lines show all 22 participant High-continuity profiles per experiment; thick lines show the gridwise median. The dashed loss reference of 3.84 is descriptive, not a calibrated confidence limit. Losses are relative to the retained unrestricted/refined optimum and displayed over 0–20. Four and 16 profiles span the entire 0–1 grid below the reference in Experiments 1 and 2. Broad or flat profiles indicate weak identification. Cohort discrimination does not imply precise individual parameter identification.

These checks are conditional on the selected model and its generating parameters. Simulation and fitting share the same model implementation, so recovery assesses estimation and discrimination within that specification rather than independently establishing its psychological mechanism. The simulation designs, optimizer records, complete tables, and additional plots are available in the public repository (see Data and Code Availability).

## Appendix C: A Test With Manipulated Sequence Volatility

The present experiments drew durations independently on every trial. To test the model under a manipulation of sequence volatility, we used the public data of Glasauer and Shi (2022). Fourteen participants reproduced durations in two conditions of 400 trials each that contained the same durations in different orders: a randomized order, and a random-walk order in which each duration was close to the preceding one (the correlation between current and previous duration was .99 in the random-walk and .02 in the randomized order).

### Models

We applied the model selected for Experiment 1, C3-S0-B2. The design has no coherence manipulation, so the C and S dimensions are inactive, and the model reduces to the three-state filter with one retention parameter *φ* and the response-level bias B2 (seven free parameters). For comparison we fitted the two-state model of Glasauer and Shi (2022) with the authors’ code (logarithmic encoding; fitted quantities *v* / *r*, *q* / *r*, and an offset), and the three-state model without the bias state (B0).

### Transfer test

Models were fitted per participant to randomized responses and then predicted the random-walk sequence with every parameter fixed; random-walk responses never entered fitting. Central tendency was one minus the reproduction–current-duration slope; serial dependence was the error–previous-duration slope. With highly correlated adjacent durations, these measures largely reflect sequence order, so the test concerns prediction of their change under fixed parameters.

### Results

The order of the sequence changed both measures strongly (Figure C1). Central tendency fell from 0.455 in the randomized to 0.096 in the random-walk condition (*t*(13) = 7.29, *p* < .001; 14 of 14 participants), and serial dependence changed sign, from 0.109 to −0.086 (*t*(13) = 5.16, *p* < .001; 13 of 14). With parameters fixed at the randomized fit, the three-state model predicted the random-walk values (Figure C1; paired comparisons with the data, *p* = .90 and .96). The two-state model predicted values closer to zero (differences from the data: *p* = .11 and .12). Scored on the held-out random-walk responses, the mean log predictive density of the three-state model was higher than that of the two-state model (difference = 0.084, 95% CI [0.030, 0.148], exact sign-flip *p* = .009; 11 of 14 participants) and than that of the model without the bias state (difference = 0.072, 95% CI [0.026, 0.127], *p* = .007; 13 of 14). The root-mean-square error of the predicted mean did not differ between models (*p* ≥ .49), so the bias state improved the predicted distribution of the responses, not their mean.

**Figure C1:**
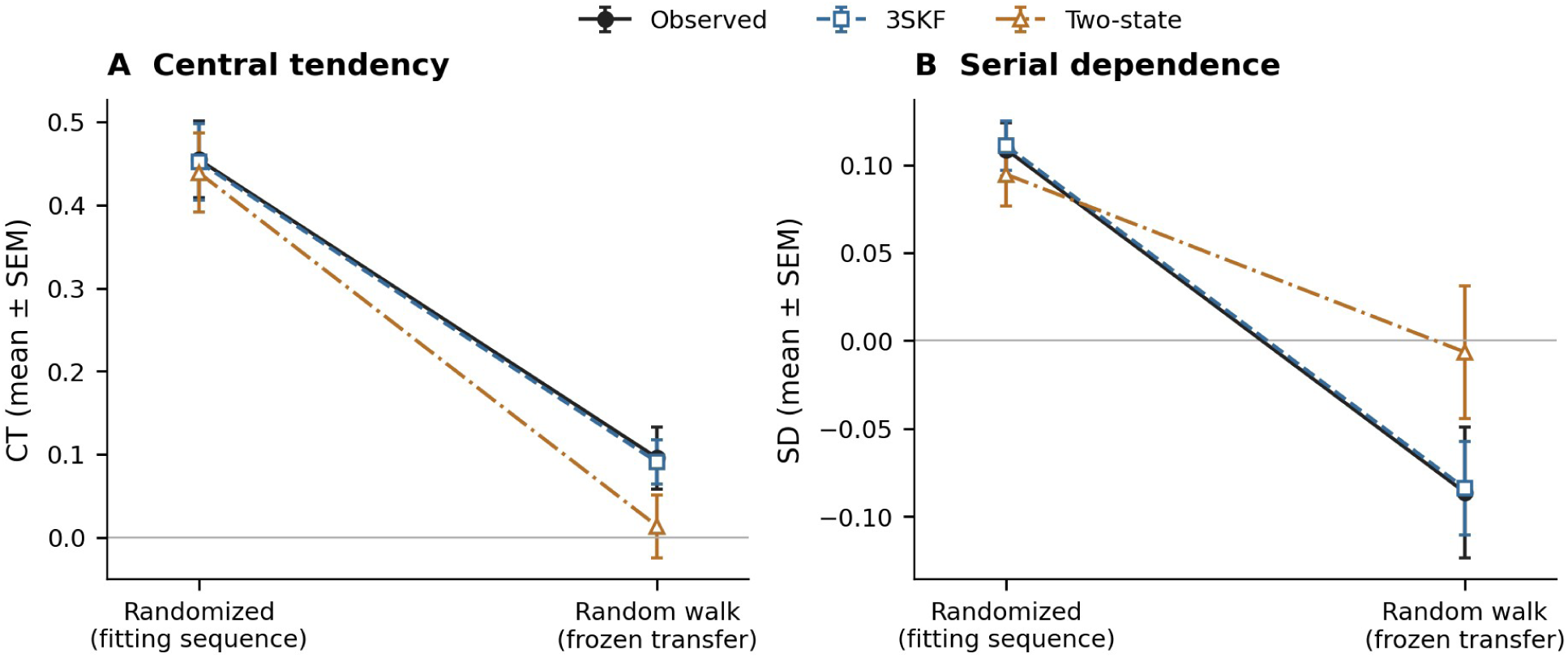
Transfer From the Randomized to the Random-Walk Sequence. *Note.* (A) Central tendency and (B) serial dependence. Observed values use filled black circles/solid lines; the three-state model uses open squares/dashed lines and the two-state model open triangles/dash-dot lines. All bars are participant SEMs, not confidence intervals. Parameters were fitted to the randomized condition and fixed before prediction of the random-walk condition. Complete participant-level summaries are available in the public repository.

### Parameters by sequence

Separate three-state fits showed no reliable sequence differences in fast/slow process noise, retention *φ*, or gains (Wilcoxon signed-rank tests, all *p* > .06). Response-noise SD alone was smaller in the random-walk order (median 0.124 versus 0.172 s, *p* = .017). The sequence effect was thus compatible with fixed observer parameters.

