## Appendix for "Precision-Weighted Updating Explains Serial Dependence Across Sensory and Contextual Transitions"

### Appendix A: Linear Mixed Models and Behavioral Analyses

We report the primary trial-level LMMs and four coherence transitions (A.1), response-error history (A.2), mean reproduction and variability (A.3), and predicted slope orderings (A.4). Complete numerical summaries and the values plotted in the figures are available in the public repository (see Data and Code Availability).

#### A.1 Duration History and Coherence Transitions

#### Primary inferential models

The primary LMMs modeled current reproduction bias as a function of centered current duration, centered previous duration, and the experiment-specific moderator. Models were fitted with restricted maximum likelihood and included participant random intercepts plus random slopes for current and previous duration:

$$\begin{aligned} E_{\mathrm{ni}} &= \beta_0 + \beta_1 T_{\mathrm{ni}} + \beta_2 T_{n-1,i} + \beta_3 M_{\mathrm{ni}} \\ &+ \beta_4 T_{\mathrm{ni}} M_{\mathrm{ni}} + \beta_5 T_{n-1,i} M_{\mathrm{ni}} + u_{0i} + u_{1i} T_{\mathrm{ni}} + u_{2i} T_{n-1,i} + \varepsilon_{\mathrm{ni}} \end{aligned} \tag{1}$$

where $\mathrm{E}_{\mathrm{ni}}$ is the reproduction bias on trial $n$ for participant $i$ (reproduced minus presented duration); $T_{\mathrm{ni}}$ and $T_{n-1,i}$ are the centered current and previous durations; $M_{\mathrm{ni}}$ is the experiment-specific moderator (current low coherence in Experiment 1; Same/Switch transition in Experiment 2); $u_{0i}, u_{1i}, u_{2i}$ are participant-level random intercepts and slopes; and $\varepsilon_{\mathrm{ni}}$ is the residual. Fixed-effect estimates for the moderator interactions are shown in Figure A1A.

These are the models underlying the main text claims. In both experiments, the current duration × moderator interaction was small and non-significant, providing no evidence that central tendency was modulated by current coherence (Experiment 1) or by Same/Switch transition status (Experiment 2). The previous duration × moderator interaction was reliable in Experiment 1, showing that the controlled serial dependence channel was selectively strengthened by low current coherence, and was not reliable in Experiment 2, where the descriptive Same > Switch pattern was weaker at the trial level.

#### Four coherence transitions

To compare the same four transitions in both experiments, a common model crossed previous and current coherence. Both durations were centered at 1.2 s, and the fixed effects were current duration × previous coherence × current coherence + previous duration × previous coherence × current coherence, with participant random intercepts and random slopes for current and previous duration (4,693 and 4,654 trials, 22 participants per experiment). Figure A1B–D shows the previous-duration slopes and their planned and post-hoc contrasts. Transitions are labeled with the previous coherence first.

**Figure A1**

*Trial-Level Duration-History Effects and Coherence-Transition Contrasts*

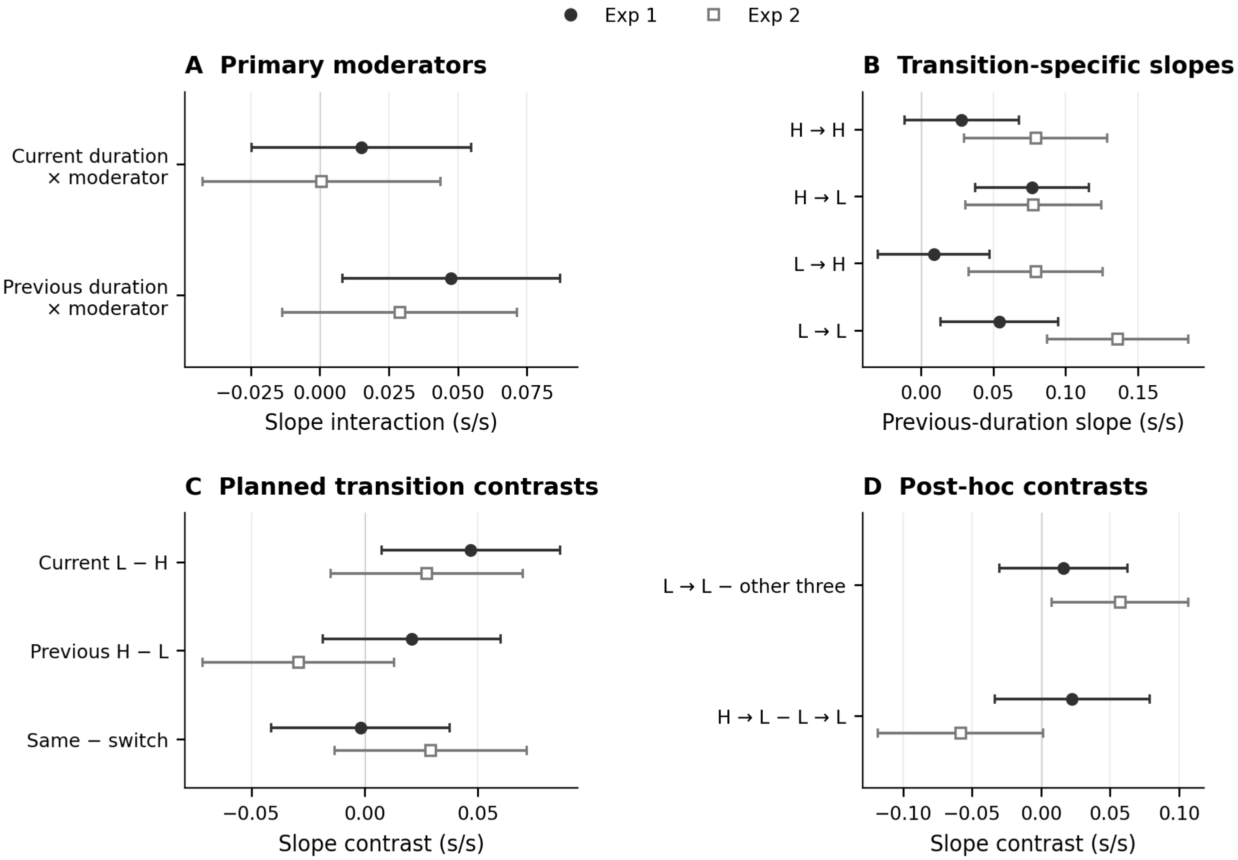


*Note.* Points and bars are the mixed-model estimates and pointwise 95% confidence intervals. Filled circles denote Experiment 1 and open squares Experiment 2. (A) Primary interaction estimates: the moderator is current Low − High coherence in Experiment 1 but Same − Switch transition in Experiment 2; these are different contrasts, not an experiment interaction. (B) Controlled previous-duration slopes; transition labels give previous then current coherence. (C) Three planned contrasts of those slopes. Their raw/Holm-adjusted $p$ pairs, in row order, are .020/.059, .298/.597, and .931/.931 in Experiment 1 and .203/.529, .176/.529, and .177/.529 in Experiment 2. (D) Four post-hoc contrasts, shown with nominal intervals and $p$ values; LL minus the other three in Experiment 2 has nominal $p$ = .024 and four-follow-up Bonferroni $p$ = .096. All intervals are unadjusted; neither interval overlap nor exclusion of zero replaces the stated correction families.

The current-coherence contrast in Experiment 1 was similar across random-effects specifications (intercept only; with either or both duration slopes) and with participant-clustered standard errors (estimates 0.046–0.047, $p$ = .006–.028). The Same − Switch estimate in Experiment 2 was positive in all specifications (0.019–0.029, $p$ = .166–.197).

### A.2 Response-Error History

The reproduction display resembled the stimulus, and each response may leave a trace of its own. To separate stimulus history from response history, we defined the response error of a preceding trial as the reproduced minus the presented duration and entered the errors of the five preceding trials together with current and previous physical duration:

$$\text{Bias}_{ij} = \beta_0 + \beta_1 T_{ij}^{\text{cur}} + \beta_2 T_{ij}^{\text{prev}} + \sum_{k=1}^{5} \gamma_k E_{ij,n\text{-}k} + u_{0j} + \varepsilon_{ij}$$

Lagged errors were constructed within participant and block on the complete trial sequence, before response exclusions (2,802 and 2,659 trials with complete histories). Response-error carryover was strongest at lag 1 and decayed across lags in both experiments (Figure A2A), consistent with a gradually decaying response-bias state. Variance inflation factors were modest (largest VIF = 1.57 and 1.46), so the lagged-error model did not suffer from problematic multicollinearity.

**Figure A2**

*Response-Error Carryover and Coherence-Dependent History Effects*

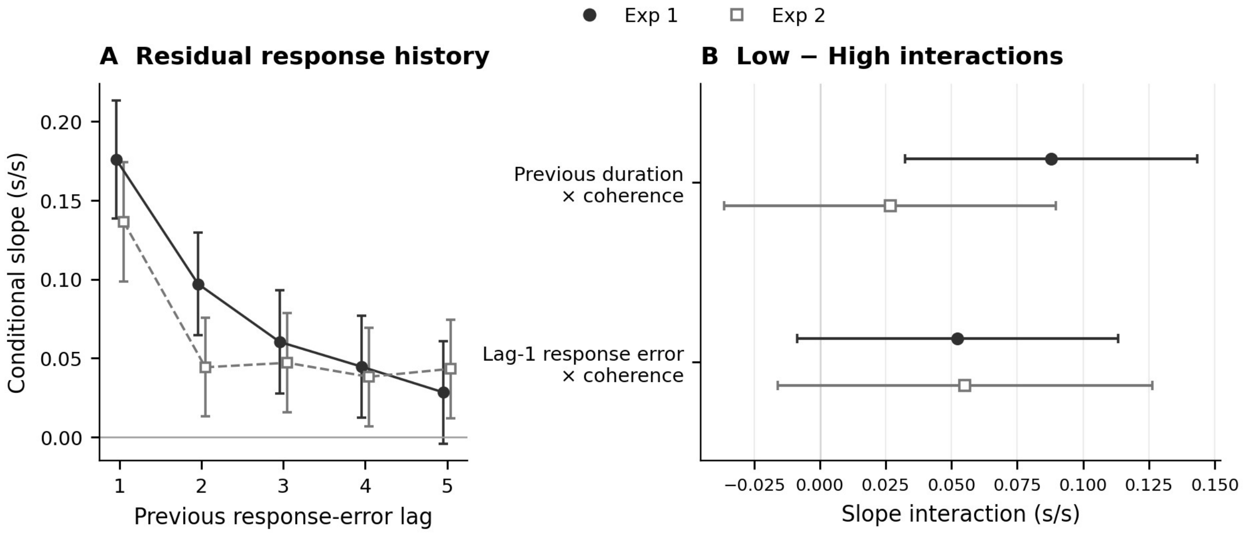


*Note.* Filled circles/solid lines denote Experiment 1; open squares/dashed lines Experiment 2. Bars are pointwise 95% confidence intervals. (A) Response-error lags 1–5 entered simultaneously, controlling current and previous duration (2,802 and 2,659 complete-history trials). (B) Low − High slope interactions from the separate coherence-interaction model, with response-error lags 2–5 as covariates. Both models include participant random intercepts.

**Figure A3**

*Mean Reproduction, Conditional Variability, and Evidence for Slope Orderings*

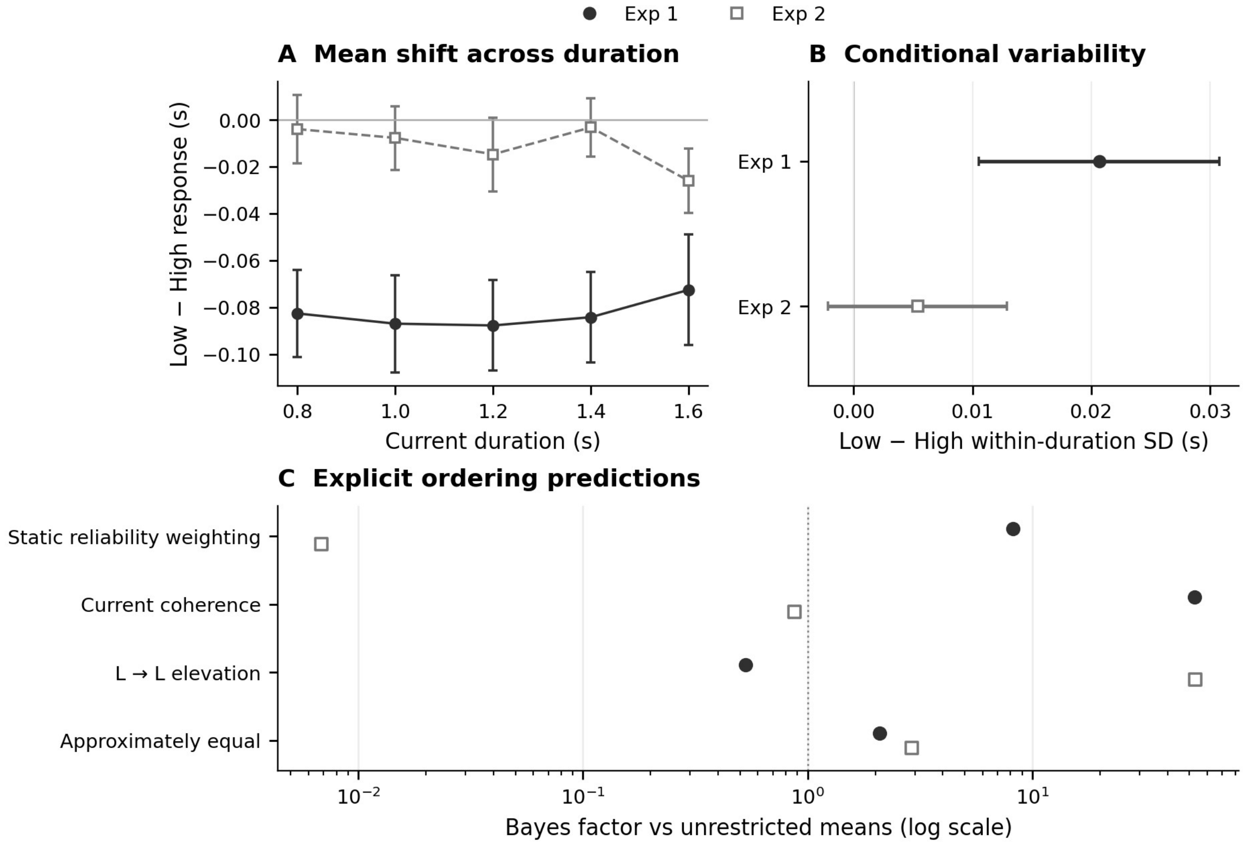


*Note.* Filled circles/solid lines denote Experiment 1; open squares/dashed lines Experiment 2. (A) Participant mean Low − High reproduction at each physical duration ± SEM; these descriptive cell means are distinct from the adjusted LMM offsets in the text. The additive-versus-proportional tests use participant slopes, not independent duration cells. (B) Within-duration reproduction-SD contrasts with paired 95% confidence intervals; this is not the residual SD of the model's predictive check. (C) Bayes factors against unrestricted slope means on a log scale, at $\tau$ = .10 and $\delta = g = .02$. The tested patterns are static reliability, HL > {LL, HH} > LH; current coherence, {HL, LL} > {HH, LH}; LL elevation, LL > {HH, LH, HL}; and approximate equality of all four transitions. Braces denote approximate equality. The reference at 1 is equal evidence against the unrestricted comparison, not a significance threshold. These are separate estimands and analyses, not a combined test.

### Appendix B: Computational Modeling Technical Details

#### B.1 The Three-State Kalman Filter

**States.** The observer's state on trial $i$ is $\boldsymbol{s}_i = \left[\hat{x}_i, \widehat{m}_i, \hat{b}_i\right]^T$, where $\hat{x}$ is the fast state (the current duration estimate), $\widehat{m}$ the slow state (the learned mean), and $\hat{b}$ the bias state. $\boldsymbol{s}_i^-$ denotes the prediction and $\boldsymbol{s}_i$ the posterior. $s_i$ is the presented duration.

**State evolution.**

$$\boldsymbol{s}_i^- = F\,\boldsymbol{s}_{i-1},\; F = \begin{bmatrix} 0 & 1 & F_{1,3} \\ 0 & 1 & 0 \\ 0 & 0 & \lambda \end{bmatrix},\; P_i^- = F\,P_{i-1}F^{\top} + Q_i$$

$$Q_i = \mathrm{diag}\left(q_1^{\mathrm{eff}}, q_2^{\mathrm{eff}}, q_3^{\mathrm{eff}}\right),\; q_j^{\mathrm{eff}} = q_j \cdot \exp\left(\alpha_{q_j}\left(1 - \mathrm{coh}_i\right)\right)$$

The modulation $\alpha_{q_j}$ is free only for the noise terms selected by the C level and is zero otherwise. The first row of $F$ gives the prediction of the fast state, $\hat{x}_i^- = \hat{m}_{i-1} + F_{1,3}\hat{b}_{i-1}$: the prediction starts from the learned mean, and the fast state of the preceding trial is not carried into the filter. The learned mean and the bias evolve as $\hat{m}_i^- = \hat{m}_{i-1}$ and $\hat{b}_i^- = \lambda\,\hat{b}_{i-1}$.

**Observation and update.** $H = [1,0,0]$. The measurement is the presented duration, $z_i = s_i$; the measurement noise $R_i$ expresses the reliability that the observer assigns to it and sets the gain.

$$R_i = \begin{cases} 1 & \mathrm{coh}_i \geq 0.5 \\ r_{\mathrm{low}} & \mathrm{coh}_i < 0.5 \end{cases}$$

$\left(r_{\mathrm{low}}\right.$ free only if the C level includes R$\left.\right)$

$$K_i = P_i^- H^{\top}\left(H P_i^- H^{\top} + R_i\right)^{-1},\; \boldsymbol{s}_i = \boldsymbol{s}_i^- + K_i\left(z_i - H\,\boldsymbol{s}_i^-\right),\; P_i = \left(I - K_i H\right)P_i^-$$

$K_i^{(x)}$ and $K_i^{(m)}$ denote the components of $K_i$ for the fast and the slow state.

**Readout.** With the fast deviation $\hat{f}_i = \hat{x}_i - \hat{m}_i$, the readout is

$$\begin{aligned} \hat{u}_i &= \hat{x}_i + \left(1 - K_i^{(x)}\right)\varphi_i \hat{f}_{i-1}, \\ \varphi_i &= \begin{cases} \varphi^L & \mathrm{coh}_i < 0.5 \\ \varphi^H & \mathrm{coh}_i \geq 0.5 \\ 0 & \text{S3 and } \mathrm{coh}_i \neq \mathrm{coh}_{i-1} \end{cases} \end{aligned}$$

This is the posterior mean under the readout prior $\tilde{x}_i^- = \hat{x}_i^- + \varphi_i \hat{f}_{i-1}$ with the gain of the filter: $\tilde{x}_i^- + K_i^{(x)}\left(z_i - \tilde{x}_i^-\right) = \hat{x}_i + \left(1 - K_i^{(x)}\right)\varphi_i \hat{f}_{i-1}$. Only $\left(\boldsymbol{s}_i, P_i\right)$ are carried to the next trial, so the readout does not feed back into the filter. With $\varphi = 0$ the readout is $\hat{x}_i$. For models without a bias in the prediction ($F_{1,3} = 0$), $\hat{x}_i^- = \hat{m}_{i-1}$ and the readout prior equals $\left(1 - \varphi_i\right)\hat{m}_{i-1} + \varphi_i \hat{x}_{i-1}$, the form used in

Equation 2 of the main text. The trace of the main text is the previous estimate $\hat{x}_{i-1}$, and the retention $\varphi_i$ is its weight in this prior.

**Response.** With the offset $d_0^{\text{eff}} = d_0 + \alpha_{d_0}(1 - \text{coh}_i)$ and response noise $\varepsilon_i \sim N(0, \sigma_e^2)$,

$$y_i = \hat{u}_i + d_0^{\text{eff}} + \varepsilon_i (B0,\ B1),\ y_i = \hat{u}_i + b_i + d_0^{\text{eff}} + \varepsilon_i (B2)$$

Under B2, $b_i = \lambda b_{i-1} + \eta_i$ with $\eta_i \sim N(0, q_3^{\text{eff}})$ is a latent response bias that the observer does not track ($F_{1,3} = 0$); the likelihood integrates over it.

#### B.2 Central Tendency and Serial Dependence in the Model

For models without a bias in the prediction (B0, B2) and without a switch effect (S0), the update is $\hat{x}_i = (1 - K_i^{(x)})\hat{m}_{i-1} + K_i^{(x)} s_i$ and $\hat{m}_i = \hat{m}_{i-1} + K_i^{(m)}(s_i - \hat{m}_{i-1})$, so that $\hat{f}_i = (K_i^{(x)} - K_i^{(m)})(s_i - \hat{m}_{i-1})$. Inserting these into the readout gives the slope of the readout on the current duration,

$$\frac{\partial \hat{u}_i}{\partial s_i} = K_i^{(x)},$$

which does not depend on $\varphi$, and the slope on the previous duration,

$$\frac{\partial \hat{u}_i}{\partial s_{i-1}} = (1 - K_i^{(x)}) \Bigg[ \underbrace{K_{i-1}^{(m)}}_{\text{learned mean}} + \underbrace{\varphi_i (K_{i-1}^{(x)} - K_{i-1}^{(m)})}_{\text{retained trace}} \Bigg].$$

Central tendency is therefore set by the gain alone. Serial dependence has two routes: the previous duration moved the learned mean, and a share $\varphi_i$ of the previous fast deviation is retained. The current coherence can change serial dependence through the factor $1 - K_i^{(x)}$, if coherence changes the gain, or through $\varphi^L \neq \varphi^H$. The gains have the closed form $K_i^{(x)} = (P_{i-1}^{(m)} + q_{1,i}^{\text{eff}}) / S_i$ and $K_i^{(m)} = P_{i-1}^{(m)} / S_i$ with $S_i = P_{i-1}^{(m)} + q_{1,i}^{\text{eff}} + R_i$, where $P^{(m)}$ is the uncertainty of the learned mean; hence $K_i^{(x)} - K_i^{(m)} = q_{1,i}^{\text{eff}} / S_i$. Under S1 the first term in the bracket is multiplied by $1 - \gamma$ on switch trials, and under B1 the term $K_{i-1}^{(b)}$ is added. These expressions agree with the implemented model to numerical precision. Figure 6C,F of the main text shows the two terms of the bracket, multiplied by $1 - K_i^{(x)}$, at the fitted parameters.

#### B.3 Model Space, Fitting, and Comparison

Every model combines one level of each of three dimensions (Table B1).

**Table B1**

*Operations Defining the Three Model Dimensions*

| Dimension | Level | Operation | Added parameters |
|---|---|---|---|
| C | C1–C15 | Coherence-dependent Low/High ratio for a non-empty subset of $\{q_1, q_2, q_3, R\}$; the subset for each C level is labeled in Figure B1 | 1–4 |
| S | S0 | No switch operation | 0 |
| S | S1 | Revert the learned mean toward $\mu_0$ on switches: $\hat{m} \leftarrow (1-\gamma)\hat{m} + \gamma\mu_0$ | 1 |
| S | S2 | Reduce the measurement gain on switches: $K \leftarrow (1-\kappa)K$ | 1 |
| S | S3 | Drop the trace of the preceding trial on switches: $\varphi_i = 0$ | 0 |
| B | B0 | No bias state | 0 |
| B | B1 | Track the bias in the duration prediction: $F_{1,3} = 1$ | 2 |
| B | B2 | Add latent bias only at response: $F_{1,3} = 0$ | 2 |

*Note.* $q_1$, $q_2$, and $q_3$ govern fast-state, learned-mean, and bias variability; $R$ governs assigned measurement reliability. $\mu_0 = 1.2$ s. Low/High ratios range from 0.1 to 10. C levels containing $q_3$ are not combined with B0, leaving 148 distinct models. All share $q_1$, $q_2$, $\varphi^L$, $\varphi^H$, $d_0$, $\alpha_{d_0}$, and $\sigma_e$; total free parameters range from 8 to 14.

**Fitting.** Models were fitted to the trial-wise presented and reproduced durations of each participant (22 per experiment) by maximum likelihood on the linear duration scale. Because the model is linear and Gaussian, the likelihood of the responses is exact; under B2 the latent response bias is integrated out with a Kalman filter over the response residuals. Trials excluded in preprocessing were presented to the observer but did not contribute to the likelihood. The learned mean was initialized at 1.2 s with a large variance, so that the first duration determines it. Each fit used 12 starting points within the parameter bounds and one additional start with $\varphi^L = \varphi^H = 0$. Bounds were set on the natural scale of each parameter: $\varphi^L$, $\varphi^H$ in [0, 1]; $\lambda$ in [0, 0.99]; the response-noise SD $\sigma_e$ in [0.01, 0.5] s; the offset $d_0$ and its Low − High difference in [−0.5, 0.5] s; $q_1$ and $q_2$ in [0, 25] in units of the high-coherence measurement noise; the SD of the response-bias drift in [0, 0.5] s; and every Low/High ratio in [0.1, 10]. Model comparison used $AIC = 2k - 2l$ (Burnham & Anderson, 2002), averaged over participants.

**Screening by simulation.** Each fitted model was simulated 20 times on each participant's trial sequence. For each statistic (central tendency and serial dependence by current coherence and by transition, and the residual SD), the model mean was compared with the data mean in units of the standard error of the data; a statistic counted as reproduced when the absolute difference was below 1.5.

**Results.** Table 1 of the main text summarizes the bias and switch dimensions. Figure B1 compares the 15 C levels in combination with the selected switch and bias levels. Differences in AIC between C levels were small, and the gain on the current duration was nearly the same under low and high coherence at every level.

**Figure B1**

*Coherence-Dependent Noise: Relative Fit and Low/High Measurement Gain*

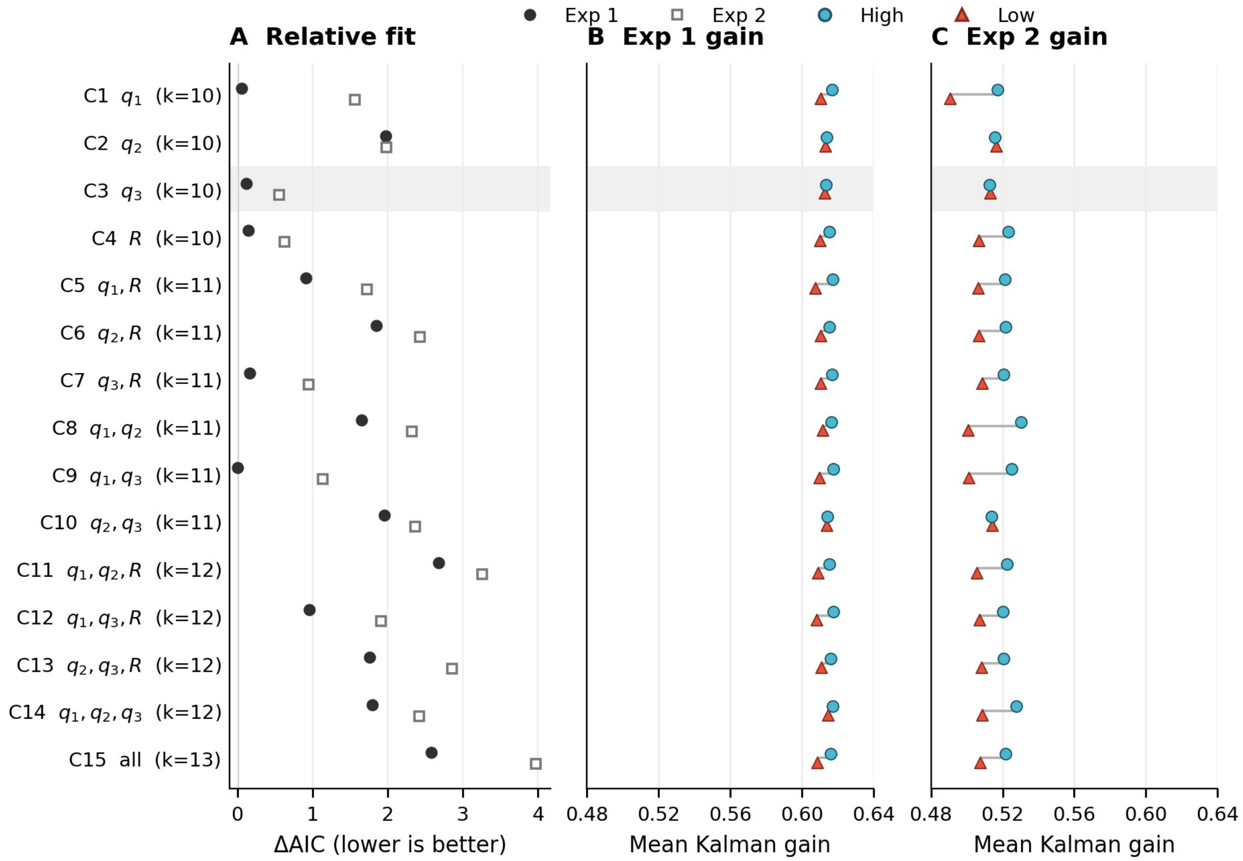


*Note.* (A) Descriptive forest-style point estimates of mean participant ΔAIC, relative to the best of the 148 models within each experiment; filled circles denote Experiment 1 and open squares Experiment 2. No confidence intervals are implied or added. Each C level is combined with S0/B2 in Experiment 1 and S3/B2 in Experiment 2. (B–C) Mean $K^{(x)}$ on current High (cyan circles) and Low (red triangles) trials, averaging within participant then across the 22 participants. Small vertical offsets distinguish overlapping markers; gray segments indicate the Low/High span, not uncertainty. The two gain panels share a scale. The shaded C3 row identifies the coherence dimension used for the retained models; the AIC minimum need not be C3. *k* is the number of free parameters.

**Parameter estimates.** Figure 6 of the main text shows the gains and the retention of the selected models. Table B2 gives their remaining parameters.

**Table B2**

*Remaining Parameter Estimates of the Selected Models*

| Parameter | Exp. 1 (C3-S0-B2): $M \pm SEM$ | Exp. 1: median | Exp. 2 (C3-S3-B2): $M \pm SEM$ | Exp. 2: median |
|---|---|---|---|---|
| $q_1$, process noise of the fast state | 2.01 ± 0.44 | 1.41 | 0.81 ± 0.16 | 0.63 |
| $q_2$, process noise of the slow state | 0.32 ± 0.24 | 0.00 | 0.17 ± 0.06 | 0.07 |
| $\sigma_b$, bias drift per trial, current Low (s) | 0.076 ± 0.014 | 0.066 | 0.047 ± 0.010 | 0.030 |
| $\sigma_b$, bias drift per trial, current High (s) | 0.061 ± 0.011 | 0.051 | 0.056 ± 0.011 | 0.044 |
| $\alpha_{q_3}$, coherence modulation of the bias drift | 0.81 ± 0.81 | 0.99 | −1.16 ± 1.00 | −2.65 |
| $\lambda$, decay of the response bias | 0.72 ± 0.06 | 0.83 | 0.78 ± 0.05 | 0.90 |
| $d_0$, response offset at coherence 1 (s) | 0.206 ± 0.034 | 0.212 | 0.001 ± 0.020 | 0.038 |
| $\Delta d_0$, Low − High response-offset difference (s) | −0.085 ± 0.017 | −0.072 | −0.012 ± 0.008 | −0.005 |
| $\sigma_e$, response-noise SD (s) | 0.134 ± 0.011 | 0.138 | 0.168 ± 0.011 | 0.171 |

*Note.* Participant means, standard errors, and medians ($N = 22$ per experiment). $q_1$ and $q_2$ are in units of the high-coherence measurement noise. $\sigma_b$ is the SD of the response-bias drift per trial at the low (30%) and the high (70%) coherence level; $\alpha_{q_3}$ is the modulation that produces the difference between the two, $q_3^{\text{eff}} = q_3 \exp\left(\alpha_{q_3}\left(1\text{-coh}\right)\right)$.

### B.4 Predictive Checks by Duration

For the predictive checks, 100 cohorts were simulated from the fitted parameters; in each cohort every participant was simulated once on their own trial sequence. Figure B2 overlays Low and High coherence within each duration-resolved check, with experiments in adjacent columns. Reproduction bias by current duration and by previous duration was reproduced in both experiments. The residual SD of the model was flat across durations. In Experiment 1 the observed residual SD increased with duration (from about 0.18 to 0.20 s under low and from 0.16 to 0.19 s under high coherence), so that the model was too high at short and too low at long durations; Experiment 2 showed no clear trend.

**Figure B2**

*Duration-Resolved Predictive Checks With Low and High Coherence Overlaid*

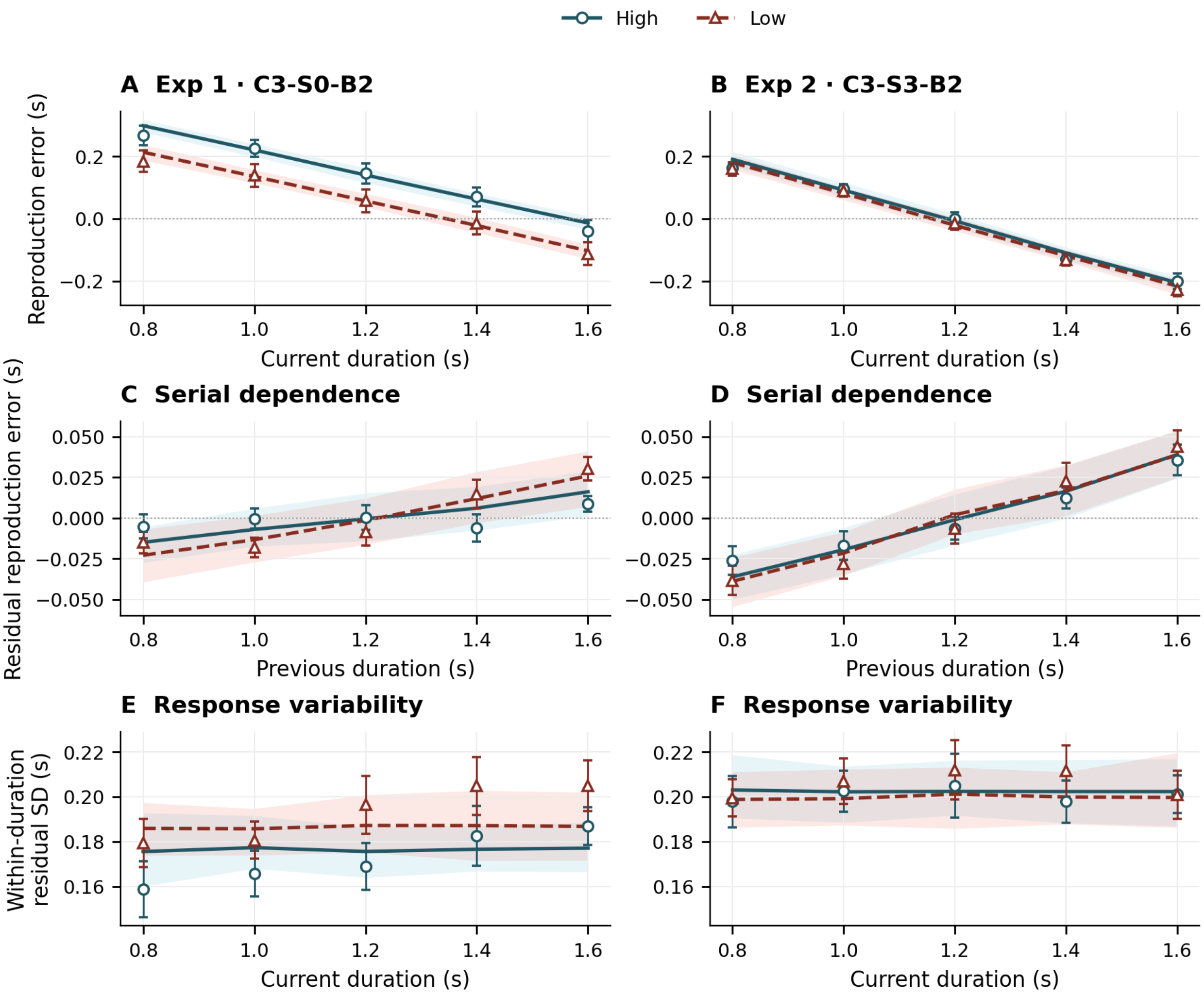


*Note.* Columns show Experiments 1 (C3-S0-B2) and 2 (C3-S3-B2); rows show bias by current duration, residual bias by previous duration, and within-duration residual SD. High coherence uses cyan circles and solid model lines; Low coherence uses red triangles and dashed lines. Open points are participant means ± SEM; model lines are means and shaded bands the central 95% range of cohort means across 100 simulated cohorts. Bands are predictive ranges, not confidence intervals for the observed means. Within each row, both experiments share a y-scale. Residual bias removes the participant/coherence/current-duration mean and is not the controlled serial-dependence slope of the LMM.

### B.5 Parameter and Coherence-Effect Recovery

To assess which fitted quantities can be estimated under the experimental design, we generated responses from the selected models, C3-S0-B2 in Experiment 1 and C3-S3-B2 in Experiment 2. Each of the 22 fitted participants per experiment generated 20 independent response datasets on their own 240-trial duration/coherence sequence, retaining the original response-validity mask. The observer processed the full sequence, including trials without a scored response. Newly drawn latent response bias and response noise varied across repetitions, and all ten free parameters were jointly re-estimated. The primary design therefore contained 880 simulated datasets. A further 400 exploratory parameter vectors sampled the fitted ranges, with continuity varying independently over 0–1; these results are available in the public repository.

The ten recovered quantities were the innovation SDs of the fast, slow, and bias states (the square roots of the process-noise variances); Low and High readout continuity (the retention parameters $\varphi^L$ and $\varphi^H$ of the main text); bias persistence; the baseline response offset and its Low − High difference; response-noise SD; and the Low/High bias innovation variance ratio. In the implementation, innovation SDs are optimized in seconds; the filter uses normalized process variances $\left(\sigma_{q_j}/\sigma_s\right)^2$, with $\sigma_s = 0.1$ fixed in these selected models. The baseline offset and bias innovation SD are defined at coherence 1, rather than at the High condition. Condition offsets are $d_H = d_0 + 0.75\,\Delta d_0$ and $d_L = d_0 + 1.75\,\Delta d_0$. Both selected C3 models have $R_H = R_L = 1$.

Initial fits used 13 starting points: the default point, 11 random points, and a zero-continuity point. Fits that did not converge or fell more than 0.01 log-likelihood below the generating parameters received 49 additional starts, without using the generating parameters as an optimization start. Original and refined outputs were retained separately, and refinements used identical simulated responses. All 880 primary selected fits converged and met the generating-likelihood check after refinement. Figure B3 distinguishes recovery from one dataset per person from recovery after averaging 20 estimates per person. Full bias, RMSE, and boundary summaries are available in the public repository.

Fast innovation SD and response offsets recovered more consistently than individual readout continuity. For High continuity, single-cohort correlations were .157 and .258 in Experiments 1 and 2, compared with .663 and .690 after averaging each participant's 20 estimates. High continuity reached its lower bound in 46.8% and 35.7% of recoveries. Thus, the parameter estimates do not support precise individual differences in High continuity. Figure B3 compares both recovery summaries for all ten fitted parameters; averaging repeated datasets must not be read as recovery from a single experiment.

**Figure B3**

*Single-Dataset Versus Repeated-Dataset Parameter Recovery*

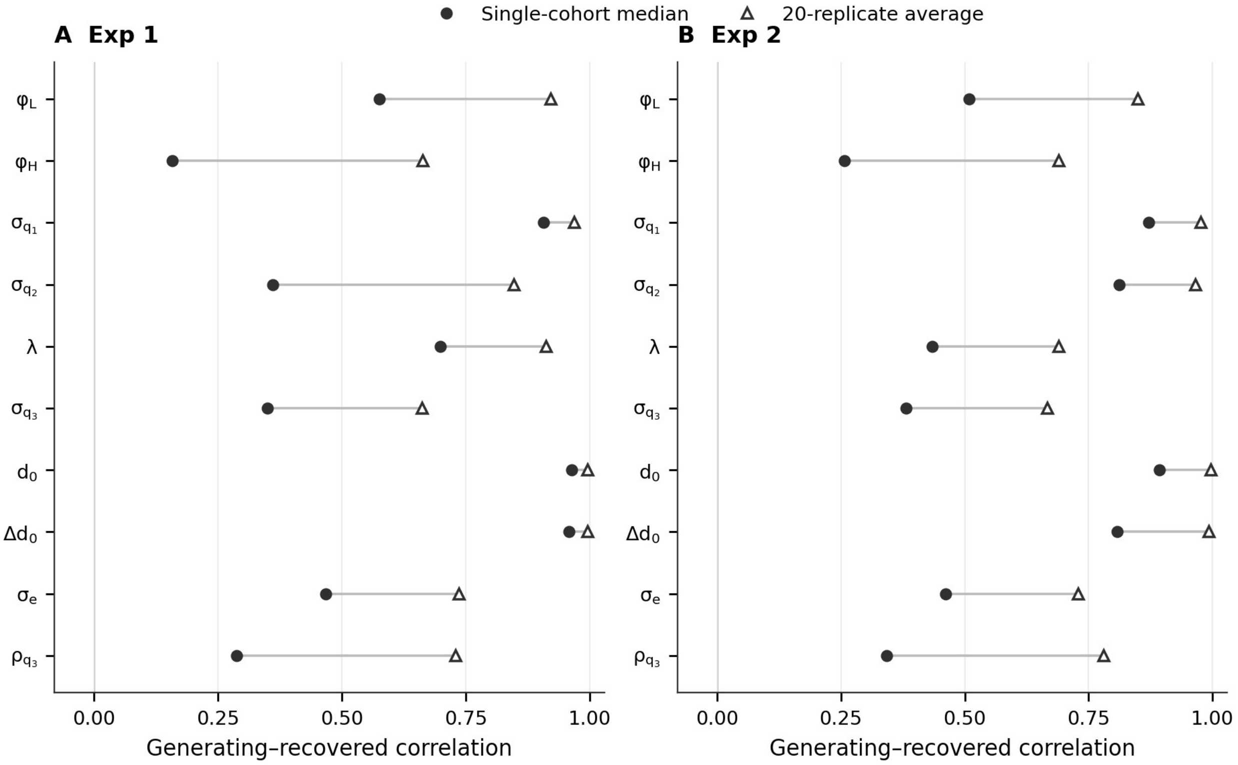


*Note.* Filled circles show the median generating–recovered correlation over 20 cohorts, each with one dataset for each of 22 participants. Open triangles show the correlation of the 22 generating values with participant estimates averaged over 20 datasets. Connecting segments distinguish these two summaries and are not confidence intervals. The two experiments share the correlation scale. $\sigma_{q_j}$ are innovation SDs; $\lambda$ is bias persistence; $d_0$ and $\Delta d_0$ are baseline and Low − High offsets; $\sigma_e$ is response noise; and $\rho_{q_3}$ is the bias-variance ratio. The weak single-dataset recovery of $\varphi^H$ remains visible rather than being obscured by the higher repeated-dataset correlations.

We also recovered the Low − High continuity difference, $\Delta\varphi = \varphi^L - \varphi^H$, and the corresponding difference in effective trace weight, $\Delta w = \overline{w}^L - \overline{w}^H$. Trial weights were computed as $w_i = \left(1 - K_i^{(x)}\right)\varphi_i$ and averaged separately over Low and High trials on the actual sequence. Under S3, switch trials contribute zero trace weight. These readout weights are distinct from the complete slope on the preceding duration, which also includes the learned-mean route (B.2). The mean generating effects were positive in both experiments, but their direction recovered more consistently in Experiment 1 (Figure B4).

**Figure B4**

*Recovery of Cohort-Mean Coherence Differences*

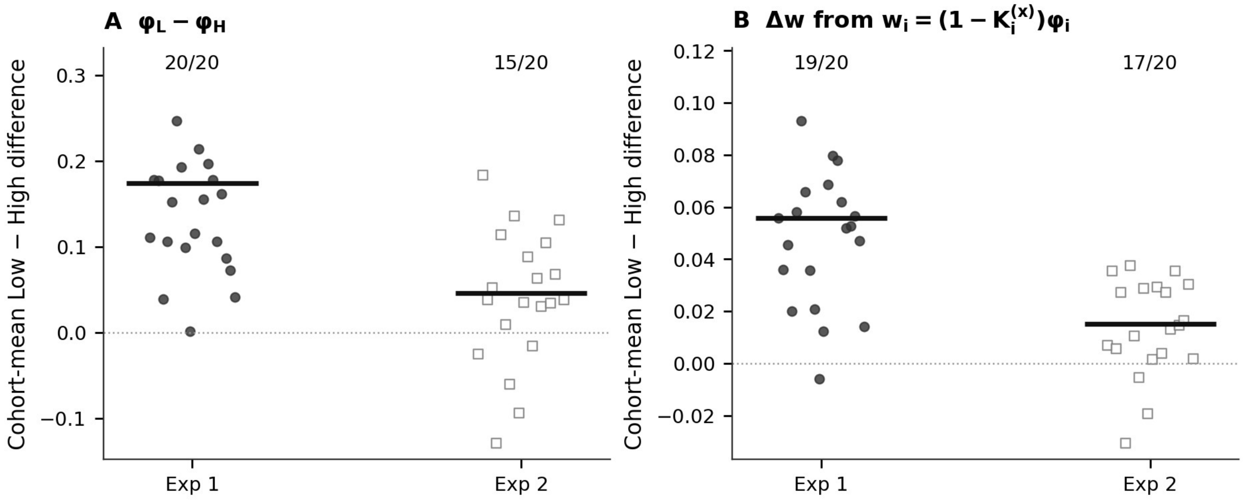


*Note.* Each point is one recovered 22-participant cohort mean, with 20 independent response cohorts per experiment; filled circles denote Experiment 1 and open squares Experiment 2. Horizontal offsets only separate repetitions. Short thick horizontal lines show the fixed generating cohort means. Panels show (A) $\Delta\varphi$ and (B) $\Delta w$, with $w_i = \left(1 - K_i^{(x)}\right)\varphi_i$ averaged within current coherence; fractions give the number of recovered cohort means with the generating sign (20/20 and 15/20 for continuity; 19/20 and 17/20 for trace weight). These are conditional recovery frequencies at the fitted generating values, not empirical-effect tests or estimates of design power.

**Figure B5**

*Cohort-Level Switch-Rule Discrimination and Individual High-Continuity Profiles*

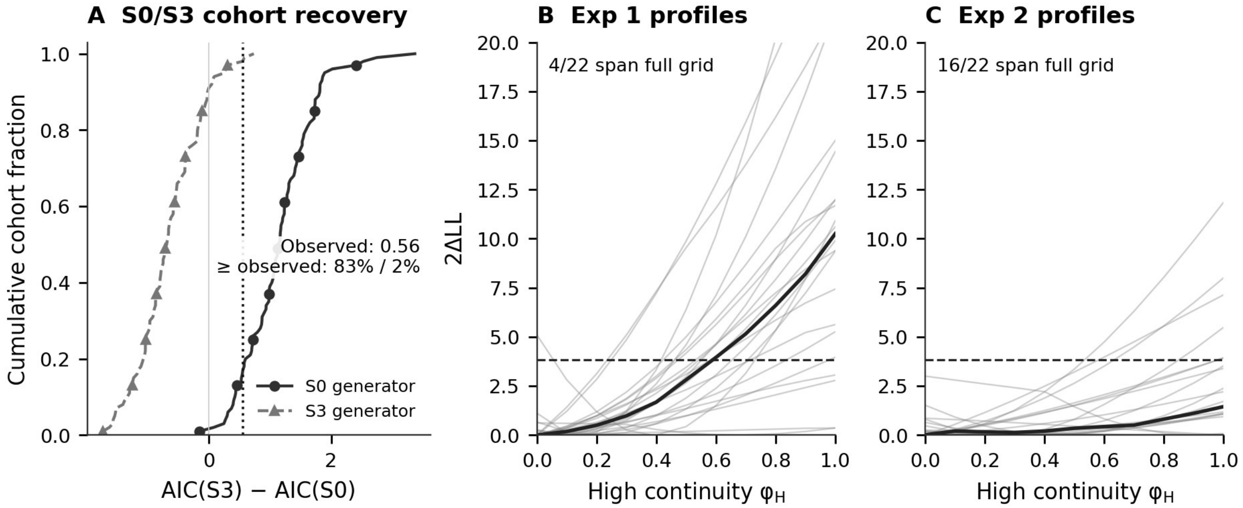


*Note.* (A) Empirical cumulative distributions of mean AIC(S3) − AIC(S0) for 100 cohorts per generating rule on Experiment 2 schedules. Solid circles denote S0-generated cohorts and dashed triangles S3-generated cohorts. Positive values favor S0; the dotted line marks the observed difference of 0.56. The 83%/2% annotation gives fractions at least that large under S0/S3 generation, not *p* values. (B–C) Thin lines show all 22 participant High-continuity profiles per experiment; thick lines show the gridwise median. The dashed loss reference of 3.84 is descriptive, not a calibrated confidence limit. Losses are relative to the retained unrestricted/refined optimum and displayed over 0–20. Four and 16 profiles span the entire 0–1 grid below the reference in Experiments 1 and 2. Broad or flat profiles indicate weak identification. Cohort discrimination does not imply precise individual parameter identification.

**Figure C1**

*Transfer From the Randomized to the Random-Walk Sequence*

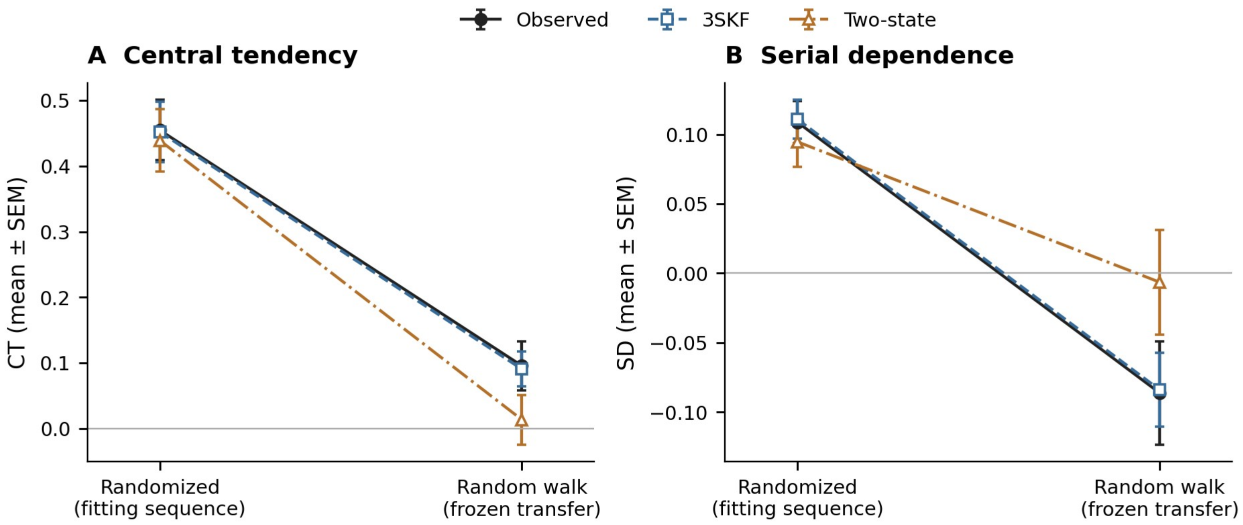


*Note.* (A) Central tendency and (B) serial dependence. Observed values use filled black circles/solid lines; the three-state model uses open squares/dashed lines and the two-state model open triangles/dash-dot lines. All bars are participant SEMs, not confidence intervals. Parameters were fitted to the randomized condition and fixed before prediction of the random-walk condition. Complete participant-level summaries are available in the public repository.

**Parameters by sequence.** Separate three-state fits showed no reliable sequence differences in fast/slow process noise, retention $\varphi$, or gains (Wilcoxon signed-rank tests, all $p >$ .06). Response-noise SD alone was smaller in the random-walk order (median 0.124 versus 0.172 s, $p = .017$). The sequence effect was thus compatible with fixed observer parameters.
